# Hierarchical Computational Modeling and Dynamic Network Analysis of Allosteric Regulation in the SARS-CoV-2 Spike Omicron Trimer Structures: Omicron Mutations Cooperate to Allosterically Control Balance of Protein Stability and Conformational Adaptability

**DOI:** 10.1101/2022.04.11.487920

**Authors:** Gennady M. Verkhivker, Steve Agajanian, Ryan Kassab, Keerthi Krishnan

**Affiliations:** Keck Center for Science and Engineering, Schmid College of Science and Technology, Chapman University, Orange, CA 92866, United States of America; Department of Biomedical and Pharmaceutical Sciences, Chapman University School of Pharmacy, Irvine, CA 92618, United States of America

## Abstract

Structural and computational studies of the Omicron spike protein in various functional states and complexes provided important insights into molecular mechanisms underlying binding, high transmissibility, and escaping immune defense. However, the regulatory roles and functional coordination of the Omicron mutations are poorly understood and often ignored in the proposed mechanisms. In this work, we explored the hypothesis that the SARS-CoV-2 spike protein can function as a robust allosterically regulated machinery in which Omicron mutational sites are dynamically coupled and form a central engine of the allosteric network that regulates the balance between conformational plasticity, protein stability, and functional adaptability. In this study, we employed coarse-grained dynamics simulations of multiple full-length SARS-CoV-2 spike Omicron trimers structures in the closed and open states with the local energetic frustration analysis and collective dynamics mapping to understand the determinants and key hotspots driving the balance of protein stability and conformational adaptability. We have found that the Omicron mutational sites at the inter-protomer regions form regulatory clusters that control functional transitions between the closed and open states. Through perturbation-based modeling of allosteric interaction networks and diffusion analysis of communications in the closed and open spike states, we quantify the allosterically regulated activation mechanism and uncover specific regulatory roles of the Omicron mutations. The network modeling demonstrated that Omicron mutations form the inter-protomer electrostatic bridges that connect local stable communities and function as allosteric switches of signal transmission. The results of this study are consistent with the experiments, revealing distinct and yet complementary role of the Omicron mutational sites as a network of hotspots that enable allosteric modulation of structural stability and conformational changes which are central for spike activation and virus transmissibility.

## Introduction

The rapidly growing body of structural, biochemical and functional studies established that the mechanism of SARS-CoV-2 infection may involve conformational transitions between distinct functional forms of the SARS-CoV-2 viral spike (S) glycoprotein.^1–9^ The S protein consists of conformationally adaptive amino (N)-terminal S1 subunit and structurally rigid carboxyl (C)-terminal S2 subunit, where S1 includes an N-terminal domain (NTD), the receptor-binding domain (RBD), and two structurally conserved subdomains SD1 and SD2 that collectively coordinate protein response to binding partners and regulate the interactions with the host cell receptor. Conformational transitions of the S protein from the closed state to the open state are exemplified by large scale movements of the RBDs that can spontaneously fluctuate between “RBD-down” and “RBD-up” positions, where binding to the host cell receptor ACE2 preferentially stabilizes the receptor-accessible “up” conformation.^1–9^

Dynamic structural changes that accompany SARS-CoV-2 S binding with the ACE2 host receptor were described in cryo-EM experiments showing a cascade of conformational transitions from a compact closed form weakened after furin cleavage to the partially open states and subsequently the ACE2-bound open form thus priming the S protein for fusion.^10, 11^ The cryo-EM and tomography examined conformational flexibility and distribution of S trimers in situ on the virion surface^12^ showing that spontaneous conformational changes and population shifts between different functional states can be maintained in different biological environments, reflecting the intrinsic properties of conformational landscapes for SARS-CoV-2 S trimers. Single-molecule Fluorescence (Förster) Resonance Energy Transfer (smFRET) studies of SARS-CoV-2 S trimer on virus particles revealed a sequence of conformational transitions from the closed state to the receptor-bound open state suggesting that mechanisms of conformational selection and receptor-induced structural adaptation can work synchronously and showing that SARS-CoV-2 neutralization can be achieved by antibodies that either directly compete with the ACE2 receptor binding or exert their effect by allosterically stabilizing the S protein in its RBD-down conformation. ^13^ Biophysical studies demonstrated that conformational dynamics and allosteric modulation of the S protein are intrinsically related, revealing long-range allosteric modulation of the RBD equilibrium, which in turn regulates exposure of the ACE2-binding site.^14, 15^ Conformational dynamics of SARS-CoV-2 S in the absence or presence of ligands was examined using smFRET imaging assays showing that ACE2 binding is controlled by the intrinsic conformational dynamics of the RBD and mechanism of conformational selection in which ACE2 captures the intrinsically accessible up conformation rather than inducing a conformational change.^14^ Moreover, this study also demonstrated that antibodies that target diverse epitopes that are distal to the RBD shift the RBD equilibrium S protein toward the up conformation, enhancing ACE2 binding, while S-D614G variant lives in an equilibrium where the RBD favors the up conformation prior to antibody binding. The recent smFRET experiments provided evidence that S protein variants D614G and E484K allosterically modulate thermodynamic preferences towards more open ACE2-accessible conformations and also can increase the kinetic accessibility and stability of the open S conformation by exhibiting slower transitions in the ACE2-accessible/bound states.^15^

The emergence of variants of concern (VOC’s) with the enhanced transmissibility and infectivity profile including D614G variant^16–19^, B.1.1.7 (alpha)^20–23^, B.1.351 (beta)^24, 25^, B.1.1.28/P.1 (gamma)^26^, B.1.1.427/B.1.429 (epsilon) variants^27, 28^, and Omicron (B.1.1.529)^29–33^ revealed complex and diverse mechanisms underlying thermodynamics and kinetic mechanisms of the S proteins, particularly showing how mutation-induced modulation of the conformational landscapes can allow for specific protein responses and structural adaptability to binding partners in different biological environments. In particular, structural and biophysical studies characterized a significant heterogeneity and plasticity of the SARS-CoV-2 S proteins including B.1.1.7 (alpha), B.1.351 (beta), P1 (gamma) and B.1.1.427/B.1.429 (epsilon) variants showing that the intrinsic conformational flexibility of the S protein variants can be controlled by mutations and determine their ability to evade host immunity and incur resistance to antibodies.^34–39^ Structural studies of the SARS-CoV-2 S protein complexes with multiple classes of potent antibodies highlighted the link between conformational plasticity and significant virus potential of spike proteins for eliciting specific binding and broad neutralization responses.^40–53^

The Omicron (B.1.1.529) variant bears 37 mutations in the S protein relative to the original SARS-CoV-2 strain with 15 mutations featured in the RBD regions. The enormous public interest and biomedical importance of this new variant led to a plethora of studies aiming to dissect the role and mechanisms underlying the diverse coverer of Omicron mutations variants.^39–43^ The latest structural investigation convincingly demonstrated that Omicron mutational diversity can induce a widespread escape from neutralizing antibody responses.^32^ According to this study, mutations S477N, Q498R, and N501Y increase ACE2 affinity by 37-fold, serving to anchor the RBD to ACE2, while allowing the RBD region freedom to develop further mutations, including those that reduce ACE2 affinity in order to evade the neutralizing antibody response.^32^ A comparison of binding affinity changes showed the higher affinity for N501Y, E484K, S477N, and Q498R, where Q498R and N501Y increased binding affinity by 26-fold and adding the S477N mutation produced a 37-fold increase in binding.^32^ Similar binding affinity patterns were observed in surface plasmon resonance (SPR) experiments of the Omicron RBD interactions with ACE2 using demonstrating tighter binding as comparison to other variants.^33^

Recent structural biology studies of the Omicron variant in various functional states and complexes with the ACE2 and several antibodies provided the initial rationale on which molecular features are responsible for high transmissibility while escaping immune defense. The crystal and cryo-EM structures of the Omicron RBD-ACE2 complex, and X-ray structure of Delta RBD-ACE2 identified the role of key residues for receptor recognition showing that mutations S477N, Q493R, Q498R and N501Y can enhance the binding affinity of RBD with ACE2 while T478K may influence ACE2 binding allosterically.^54^ The cryo-EM structural analysis of the Omicron variant in the unbound form and in the complex with ACE2 revealed that the mutations Q493R, G496S, Q498R and N501Y can enhance the ACE2 binding strength and offset some loss of affinity caused by other RBD mutations.^55^ Biophysical experiments studies suggested that the Omicron variant may have evolved to mediate diverse neutralization escape via multiple mutations while enhancing binding affinity with ACE2 using mutational changes in several key energy hotspots.^55^ The cryo-EM analysis of the Omicron S trimers showed that Omicron substitutions in the S2 subunit can strengthen the interaction network between neighboring protomers and subunits in the closed state which may be important for regulatory functions of the S protein.^56^ At the same time, another group of Omicron RBM mutations Q493R, and Q498R, G496S and Y505H can mediate the improved binding with the ACE2 receptor.^56^ These structural insights suggested that Omicron mutations can be important in modulating both local and long-range interactions in the S protein, providing a diverse range of modulation functions through a carefully orchestrated cooperation between mutated positions. The cryo-EM structures of the SARS-CoV-2 Omicron S ectodomain trimer bound to S309 and S2L20 antibodies and the X-ray crystal structure of the Omicron RBD in complex with human ACE2 further elucidated structural basis of Omicron-induced immune evasion and receptor engagement mechanisms^57^. This study confirmed that the favorable interactions mediated by S477N, Q493R, Q496S, Q498R, and N501Y positions can offset a partial loss of polar interactions caused by K417N and E484A mutations, indicating that the latter two mutations may have evolved to regulate immune evasion rather than binding. ^57^ Structural and biochemical characterizations of the S Omicron trimer demonstrated that ACE2 binding can stabilize the S protein in the 1 RBD-up state through the formation of favorable inter-protomer RBD-RBD contacts and the improved ACE2-binding interactions that collectively contribute to the 6-9 fold increased affinity of the Omicron S protein as compared to the wild-type S trimer.^58^ Structural diversity of the SARS-CoV-2 Omicron S protein was revealed in a series of cryo-EM structures of the Omicron and Delta variants in different functional states, showing the reduced S1 variability with immune-evasive RBD substitutions stabilizing the RBD-RBD interfaces in the 3-RBD-down closed S structure.^59^ While most of the VOC’s are implicated in the enhanced viral transmissibility through mutation-induced stabilization of the open S states, this study established that the Omicron variant can lead to the increased thermodynamic stabilization of the RBD-down closed state, which was proposed to promote immune evasion by occluding immunogenic sites.^59^ To explain the high transmissibility of the Omicron variant which requires acquisition of the open S states to engage receptor, structural analysis pointed to unusual rearrangements in the N2R linker that connects the NTD and RBD and primed to transition to the open state. However, the detailed thermodynamic and kinetic analysis of the dynamic equilibrium between the closed and open Omicron states is still lacking raising important questions concerning the mechanistic roles of Omicron mutations in controlling protein stability and dynamic changes in the S protein. The cryo-EM structures of the Omicron S-trimer solved at serological and endosomal pH were consistent with other structural studies and thermal stability assays verified that the Omicron S-trimer was more stable than the wild-type and Delta variants.^60^ Another structural analysis of the binding interface with ACE2 confirmed that T478K, Q493R, G496S, and Q498R could strengthen binding of the S Omicron to ACE2, with the N501Y mutation alone improves the binding affinity by 6-fold.^61^ The cryo-EM study of the full-length S protein of the Omicron variant analyzed binding and antigenic properties of the Omicron S trimer by bio-layer interferometry (BLI) confirming the improved binding due to N501Y, Q493R and Q498R mutations.^62^ Several other recent cryo-EM structural studies supported the proposed mechanism in which Omicron mutations can stabilize the S trimer in the closed state, allowing for receptor binding while concealing epitopes that are the targets of neutralizing antibodies.^63^ The studies reaffirmed that Omicron mutations S477N, T478K and E484A in the flexible RBM region together with K417N account for only a moderate loss of the ACE2 binding while enhancing the neutralization escape potential of the Omicron variant from highly potent antibodies that target the RBM epitopes.

Structural basis for potent antibody neutralization of the Omicron variant was examined in an illuminating investigation in which multiple cryo-EM structures of antibodies bound to the S Omicron protein were revealed.^64^ This study provided a systematic structural analysis of several major classes of antibodies revealing potential mechanisms of antibody neutralization, escape and retained potency. This study showed that in the context of S trimers, Omicron mutations provide only a moderate alteration in binding affinity to ACE2, showing that affinity improving changes are used to compensate for mutations required for immune escape. This structural study claimed that variant evolution may be driven not by the optimization of ACE2 binding but primarily by immune pressure.^64^ The cryo-EM studies identified recently two highly conserved regions on the Omicron variant RBD, including a part of the lateral surface and the cryptic site inside the trimeric interface, recognized by broadly neutralizing antibodies.^65^ A bispecific single-domain antibody was designed to simultaneously and synergistically bind these two regions as revealed by cryo-EM structures and showing that this bispecific antibody can exhibit significant neutralization breadth and therapeutic efficacy in mouse models of SARS-CoV-2 infections. ^65^ Computer simulations provided important atomistic and mechanistic insights into understanding of the dynamics and function of SARS-CoV-2 glycoproteins. Molecular dynamics (MD) simulations of the SARS-CoV-2 S proteins and mutants detailed conformational changes and diversity of ensembles, demonstrating enhanced functional and structural plasticity of the S proteins.^66–77^ Using distributed computing MD simulations of the viral proteome observe dramatic opening of the S protein complex, predicting the existence of several cryptic epitopes in the S protein.^70^ The free energy landscapes of the functional spike states combined with nudged elastic pathway optimization mapping of the conformational transitions revealed transient allosteric pockets at the hinge regions located near the D614 position.^72^ All-atom MD simulations of the fully glycosylated S-protein in solution and targeted MD simulations of conformational changes between open and closed forms revealed the key electrostatic interdomain interactions mediating protein stability and kinetics of functional spike states.^76^ Using the generalized replica exchange MD simulations with solute tempering of selected surface charged residues the conformational landscapes of the full-length S protein trimers were investigated.^77^ The intrinsic flexibility of S-protein observed in these enhanced sampling simulations suggested driving forces of conformational transitions between functional states and unveiled previously unknown cryptic pockets in the meta-stable intermediate states that may be employed in the drug design.^77^

Structural and computational analysis analyzed the interactions with ACE2 between the Omicron RBD and wild-type RBDs in their binding to the ACE2 receptor.^78–80^ Steered molecular dynamics (SMD) simulations combined with microscale thermophoresis showed the enhanced RBD-ACE2 interactions of N501Y, Q493R, and T478K mutational sites and revealed that the Omicron RBD exhibits 5-fold higher binding affinity to ACE2 compared to the native RBD.^81^ Computer simulations of the S Omicron RBD structures and complexes with ACE2 detailed the molecular determinants of the enhanced interfacial interactions mediated by the Omicron mutational sites.^82, 83^ Recent insightful evolutionary analysis^84^ suggested that the mechanisms underlying emergence of the Omicron variant involves multiple fitness trade-offs including balancing between immune escape^85–87^ and binding affinity for ACE2 proteins^33, 55, 56^ and between the requirements to switch to the RBD-up configuration for ACE2 recognition and the increased stabilization of the closed form that prevents binding of neutralizing antibodies.^59–88^

Despite of the rapidly growing body of computational studies of the S Omicron proteins the specific regulatory roles of individual Omicron mutations remain unclear, and the potential mechanism underlying a cooperative cross-talk between mutational sites has not been examined at the atomistic level. While the existing structural and computational studies focused solely on examining the effects of Omicron mutations in the S1-RBD regions involved in ACE2 binding, functional roles of the numerous S2 Omicron sites are poorly understood and often ignored in the proposed mechanisms of the Omicron-induced effects of virus infectivity and enhanced transmissibility. In this work, we further expand on our previous studies showing that the SARS-CoV-2 S protein can function as a robust and efficient allosteric regulatory engine that can exploit the intrinsic plasticity of functional regions and versatility of allosteric hotspots to modulate specific regulatory and binding functions.^89–94^ We propose that Omicron mutational sites can act in a cooperative manner and through allosteric-based mechanism control balance and trade-offs between conformational plasticity, protein stability, and functional adaptability. To overcome limitations associated with the cost of atomistic simulations for the multiple full-length SARS-CoV-2 S Omicron structures, we employed a range of methods based on efficient coarse-grained (CG) simulations and high resolution atomistic graph models combined with dynamic network analysis. We combined CG Brownian dynamics simulations of multiple full-length SARS-CoV-2 S Omicron trimers structures in the closed and open states with the local frustration analysis of conformational ensembles, mutational analysis of protein stability and extensive dynamic modeling of allosteric interaction networks. These tools are computationally efficient and can provide insights into the global dynamic and allosteric signatures of the S Omicron trimer structures. By combining CG simulations with the local frustration analysis of protein residues and modeling of collective motions, we show that the clusters of Omicron mutational sites at the inter-protomer regions of the S2 subunit are involved in interactions with the inter-protomer and inter-domain hinge residues, forming local regulatory clusters that control functional movements and transitions between the closed and open states. This study shows that Omicron sites collectively form a tightly integrated network in which S2 Omicron positions function as stable regulatory hotspots that allosterically communicate with the flexible S1-RBD Omicron positions and control a range of functional RBD movements required for regulatory and binding responses to antibody binding without compromising binding affinity with ACE2. Through perturbation-based scanning of allosteric interaction networks and communications in the closed and open S Omicron states, we quantify the specific functional role and long-range communication between sites targeted by Omicron mutations. The results of this study show that the Omicron mutations can cooperatively interact to adaptively modulate the regulatory and binding functions of the S protein. The proposed strategy can provide am important complementary framework to more rigorous and detailed mesoscale simulations of SARS-CoV-2 S proteins.

## Materials and Methods

### Structure Preparation and Refinement

All structures were obtained from the Protein Data Bank.^95, 96^ Hydrogen atoms and missing residues were initially added and assigned according to the WHATIF program web interface.^97, 98^ The structures were further pre-processed through the Protein Preparation Wizard (Schrödinger, LLC, New York, NY) and included the check of bond order, assignment and adjustment of ionization states, formation of assignment of partial charges as well as additional check for possible missing atoms and side chains that were not assigned in the initial processing with the WHATIF program. The missing loops in the cryo-EM structures were reconstructed using template-based loop prediction approaches ModLoop^99^ and ArchPRED^100^ and further confirmed by FALC (Fragment Assembly and Loop Closure) program.^101^ The side chain rotamers were refined and optimized by SCWRL4 tool.^102^ The protein structures were then optimized using atomic-level energy minimization using 3Drefine method.^103^

In addition to the experimentally resolved glycan residues present in the structures of studied SARS-CoV-2 S-RBD complexes, the glycosylated microenvironment was mimicked by using the structurally resolved glycan conformations for 16 out of 22 most occupied N-glycans (N122, N165, N234, N282, N331, N343, N603, N616, N657, N709, N717, N801, N1074, N1098, N1134, N1158) as determined in the cryo-EM structures of the SARS-CoV-2 spike S trimer in the closed state (pdb id 6VXX) and open state (pdb id 6VYB) and the cryo-EM structure SARS-CoV-2 spike trimer (K986P/V987P) in the open state (pdb id 6VSB).

### Brownian Dynamics Simulations

Coarse-grained Brownian dynamics (BD) simulations have been conducted using the ProPHet (Probing Protein Heterogeneity) approach and program.^104–107^ BD simulations are based on a high resolution CG protein representation^108^ of the SARS-CoV-2 S Omicron trimer structures that can distinguish different residues. In this model, each amino acid is represented by one pseudo-atom at the Cα position, and two pseudo-atoms for large residues. The interactions between the pseudo-atoms are treated according to the standard elastic network model (ENM) in which the pseudo-atoms within the cut-off parameter, *R*_c_ = 9 Å are joined by Gaussian springs with the identical spring constants of *γ* = 0.42 N m^−1^ (0.6 kcal mol^−1^ Å^−2^. The simulations use an implicit solvent representation via the diffusion and random displacement terms and hydrodynamic interactions through the diffusion tensor using the Ermak-McCammon equation of motions and hydrodynamic interactions as described in the original pioneering studies that introduced Brownian dynamics for simulations of proteins.^109, 110^ The stability of the SARS-CoV-2 S Omicron trimers was monitored in multiple simulations with different time steps and running times as prescribed in the original studies. ^106, 107^ We adopted Δ*t* = 5 fs as a time step for simulations and performed 100 independent BD simulations for each system using 100,000 BD steps at a temperature of 300 K.

### Distance Fluctuations Stability and Communication Analysis

We employed distance fluctuation analysis of the simulation trajectories to compute residue-based stability profiles. The fluctuations of the mean distance between each pseudo-atom belonging to a given amino acid and the pseudo-atoms belonging to the remaining protein residues were computed. The fluctuations of the mean distance between a given residue and all other residues in the ensemble were converted into distance fluctuation stability indexes that measure the energy cost of the residue deformation during simulations.^104–107^ The distance fluctuation stability index for each residue is calculated by averaging the distances between the residues over the simulation trajectory using the following expression:

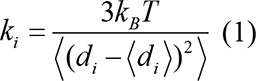

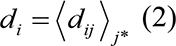

d*_ij_* is the instantaneous distance between residue *i* and residue *j*, *k_B_* is the Boltzmann constant, *T* =300K. 〈 〉 denotes an average taken over the MD simulation trajectory and 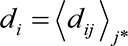 is the average distance from residue *i* to all other atoms *j* in the protein (the sum over *j*_*_ implies the exclusion of the atoms that belong to the residue *i*). The interactions between the *C_α_* atom of residue *i* and the *C_α_* atom of the neighboring residues *i* −1 and *i* +1 are excluded in the calculation since the corresponding distances are nearly constant. The inverse of these fluctuations yields an effective force constant *k_i_* that describes the ease of moving an atom with respect to the protein structure. The dynamically correlated residues whose effective distances fluctuate with low or moderate intensity are expected to communicate over long distances with the higher efficiency than the residues that experience large fluctuations. Our previous studies showed that residues with high value of these indexes often serve as structurally stable centers and regulators of allosteric signals, whereas small values of the distance fluctuation stability index are typically indicative of highly dynamic fluctuating sites.^89, 90^ The structurally stable and densely interconnected residues as well as moderately flexible residues that serve as a source or sink of allosteric signals could feature high value of these indexes. In this approach, the residue-based distance fluctuation stability index is employed as a metric for an assessment of both structural stability and allosteric propensities of the SARS-CoV-2 S residues.

### Protein Stability Calculations

A systematic alanine scanning of the SARS-CoV-2 S Omicron inter-protomer interface residues was performed using FoldX approach.^111, 112^ We utilized a graphical user interface for the FoldX calculations that was implemented as a plugin for the YASARA molecular graphics suite package.^112^ If a free energy change between a mutant and the wild type (WT) proteins ΔΔG= ΔG (MT)-ΔG (WT) > 0, the mutation is destabilizing, while when ΔΔG <0 the respective mutation is stabilizing. We computed the average ΔΔG values using multiple samples (∼ 500) from the equilibrium ensembles using a modified FoldX protocol.^113, 114^

### Dynamic-Based Modeling of Residue Interaction Network and Community Analysis

A graph-based representation of protein structures^115, 116^ is used to represent residues as network nodes and the inter-residue edges to describe non-covalent residue interactions. The network edges that define residue connectivity are based on non-covalent interactions between residue side-chains that define the interaction strength *I_ij_* according to the following expression used in the original studies.^115, 116^

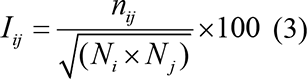

Where *n_ij_* is number of distinct atom pairs between the side chains of amino acid residues *i* and *j* that lie within a distance of 4.5 Å. *N_i_* and *N_j_* are the normalization factors for residues *i* and *j*. We constructed the residue interaction networks using both dynamic correlations^117^ and coevolutionary residue couplings^118^ that yield robust network signatures of long-range couplings and communications. The details of this model were described in our previous studies.^90–92^ In our model, the dynamics of the system is reduced to a network of correlated local motions, and signal propagation is described as an information exchange through the network.

The edge lengths in the network are obtained using the generalized correlation coefficients ***R_MI_***(***X_i_***, ***X_j_***) associated with the dynamic correlation and mutual information shared by each pair of residues. The length (i.e. weight) 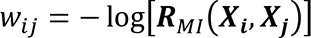 of the edge that connects nodes *i* and *j* is defined as the element of a matrix measuring the generalized correlation coefficient ***R_MI_***(***X_i_***, ***X_j_***) as between residue fluctuations in structural and coevolutionary dimensions. Network edges were weighted for residue pairs with ***R_MI_***(***X_i_***, ***X_j_***)> 0.5 in at least one independent simulation. The matrix of communication distances is obtained using generalized correlation^119^ between composite variables describing both dynamic positions of residues and coevolutionary mutual information between residues. As a result, the weighted graph model defines a residue interaction network that favors a global flow of information through edges between residues associated with dynamics correlations and coevolutionary dependencies. The ensemble of shortest paths is determined from matrix of communication distances by the Floyd-Warshall algorithm.^120^ Network graph calculations were performed using the python package NetworkX.^121^

The betweenness of residue *i* is defined as the sum of the fraction of shortest paths between all pairs of residues that pass through residue *i*:

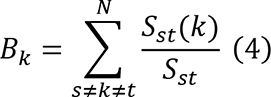

where *S*_*st*_ denotes the number of shortest geodesics paths connecting s and *t,* and *S*_*st*_ (*k*) is the number of shortest paths between residues *s* and *t* passing through the node *k*.

Using the constructed dynamic interaction networks, we performed a community analysis. A community is formed by a set of residue nodes with the stronger connection between the nodes of the community as compared to the nodes from other communities. We applied the dynamic and coevolutionary correlation to measure the interconnection between nodes and the resulting communities are composed of residues that are evolutionary coupled and move in a concerted fashion. The Girvan-Newman algorithm^122^ is used to identify local communities. In this approach, edge centrality (also termed as edge betweenness) is defined as the ratio of all the shortest paths passing through a particular edge to the total number of shortest paths in the network. The algorithm iteratively removes edges with the highest betweenness and recalculates the betweenness of the remaining edges until all edges have been removed. The modularity parameter *Q* is used to determine the goodness of the community structure. The best division occurs when nodes within a community are highly intra-connected and, at the same time, different communities are poorly inter-connected. We used a modified version of the Girvan-Newman method that was documented in detail in our recent studies.^123^ To examine network properties of the SARS-CoV-2 S Omicron states, we also computed the hierarchical community centrality. To define the hierarchical community centrality profiles, we used representative community “leaders” which are residues that are connected not only to the members of the local community but also to neighboring residues from other communities. For defining community leader nodes, we follow the Leader-Follower algorithm, in which a community is defined as a clique and is characterized by the presence of a leader and at least one ‘loyal follower’.^124^ Community leaders are defined as nodes that (a) are connected not only to members of the local community but also have neighbors outside of the community; and (b) whose distance to other nodes in the network is less than the neighbors in their respective communities. Loyal follower in a community is defined as a residue node that only has neighbors within this single community. In this model allosteric communications can be transmitted between local clusters of mega-nodes linked via inter-modular regulatory bridges. The mega-nodes of the higher hierarchical layer aggregate local communities from the lower level where nodes are simply represented by the individual residues.

### Network-Based Markov Transient Analysis

A node-based Markov Transient analysis of a random walk on the network graph was used to model kinetics of signal propagation between remote protein sits at the atomistic level. The method has successfully identified allosteric hotspots and pathways in protein systems.^125–127^ The contribution of each atom in the communication pathway between the active site and all other sites in a protein or protein complex is measured by the characteristic transient time *t*1/2, which is the number of time steps in which the probability of a random walker to be at node *i* reaches the stationary distribution value.^125, 126^ This provides a kinetic metric that estimates the time required for perturbations originating from the source site to diffuse into the rest of the protein by a random walk on the residue interaction graph. To obtain the transient time *t*_1/2_ for each residue, we take the average *t*_1/2_ over all atoms of the respective residue.

## Results and Discussion

### CG-BD Simulations and Distance Fluctuation Analysis Reveal Conformational Flexibility Patterns of the SARS-CoV-2 S Omicron states

We employed a series of independent CG-BD simulations of the multiple SARS-CoV-2 S Omicron trimer structures in the closed and open forms (Figure 1) to obtain the equilibrium dynamics profiles. Multiple cryo-EM structures of the S Omicron trimers were used in simulations (Figure 1, Table 1) allowing for a comparative analysis of protein dynamics in the distinct S Omicron states, providing the assessment of the distributions of rigid and flexible protein regions, and quantifying the effect of mutations on the mobility signatures at the Omicron positions (Figure 2).

**Figure 1.**
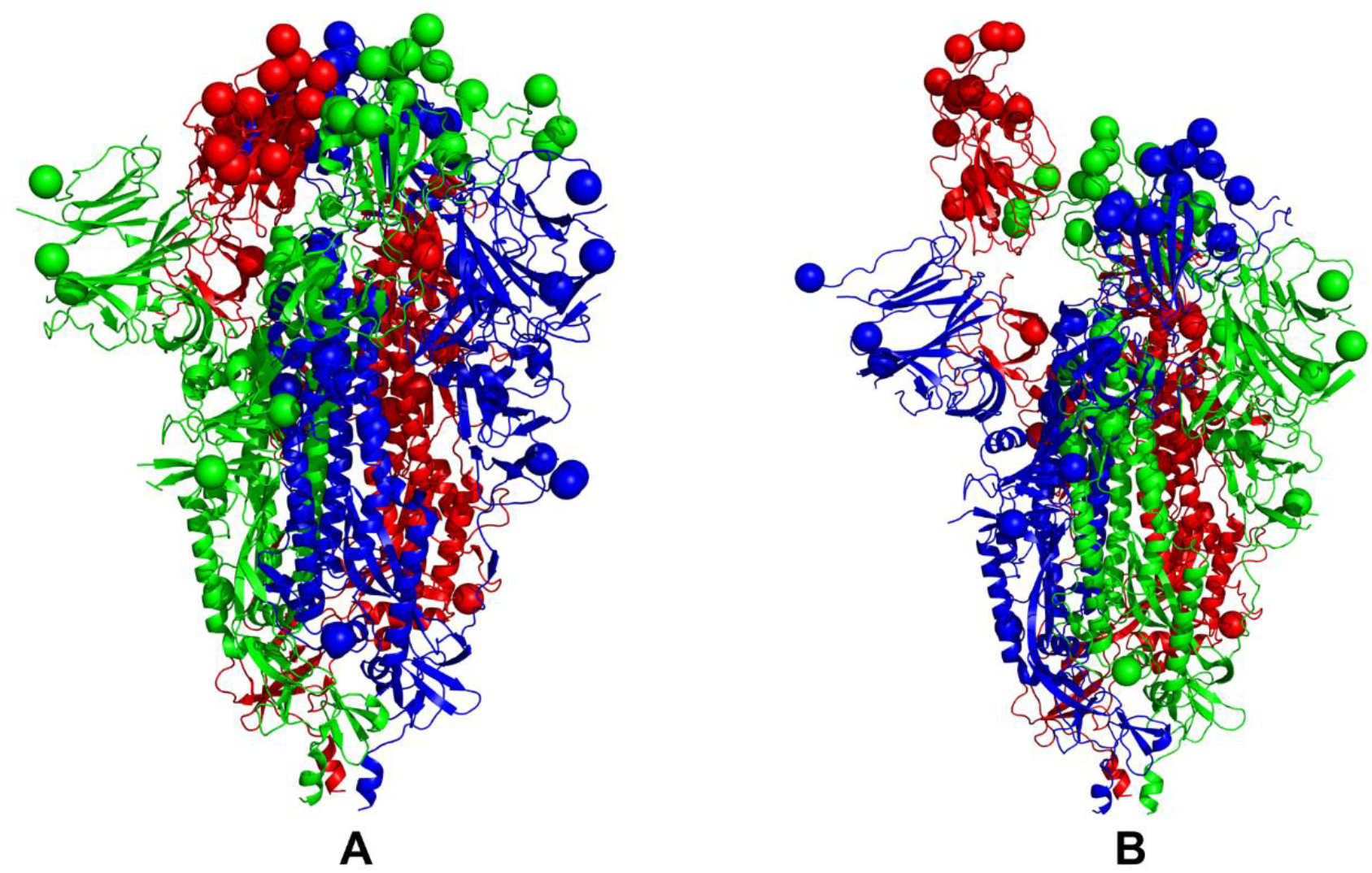
Structural organization of the SARS-CoV-2 S Omicron structures. (A) A general overview of the SARS-CoV-2 S Omicron variant in the 3RBD-down closed form (pdb id 7TF8,7WK2,7TNW) examined in this study. For clarity the cryo-EM structure of the closed S Omicron trimer, pdb id 7TF8 is shown. The protomer A is shown in green ribbons, protomer B is in red ribbons, and protomer C is in blue ribbons. The positions of Omicron mutations A67V, T95I, G142D, G339D, S371L, S373P, S375F, K417N, N440K, G446S, S477N, T478K, E484A, Q493R, G496S, Q498R, N501Y, Y505H, T547K, D614G, H655Y, N679K, P681H, N764K, D796Y, N856K, Q954H, N969K, and L981F are shown for each protomer in spheres colored according to the respective protomer. (B) A general overview of the SARS-CoV-2 S Omicron variant in the 1 RBD-up open form (pdb id 7TF8,7WK2,7TNW) examined in this study. Only the cryo-EM structure of the open S Omicron trimer, pdb id 7TEI is shown. The positions of Omicron mutations are shown in spheres colored according to the respective protomer.

**Figure 2.**
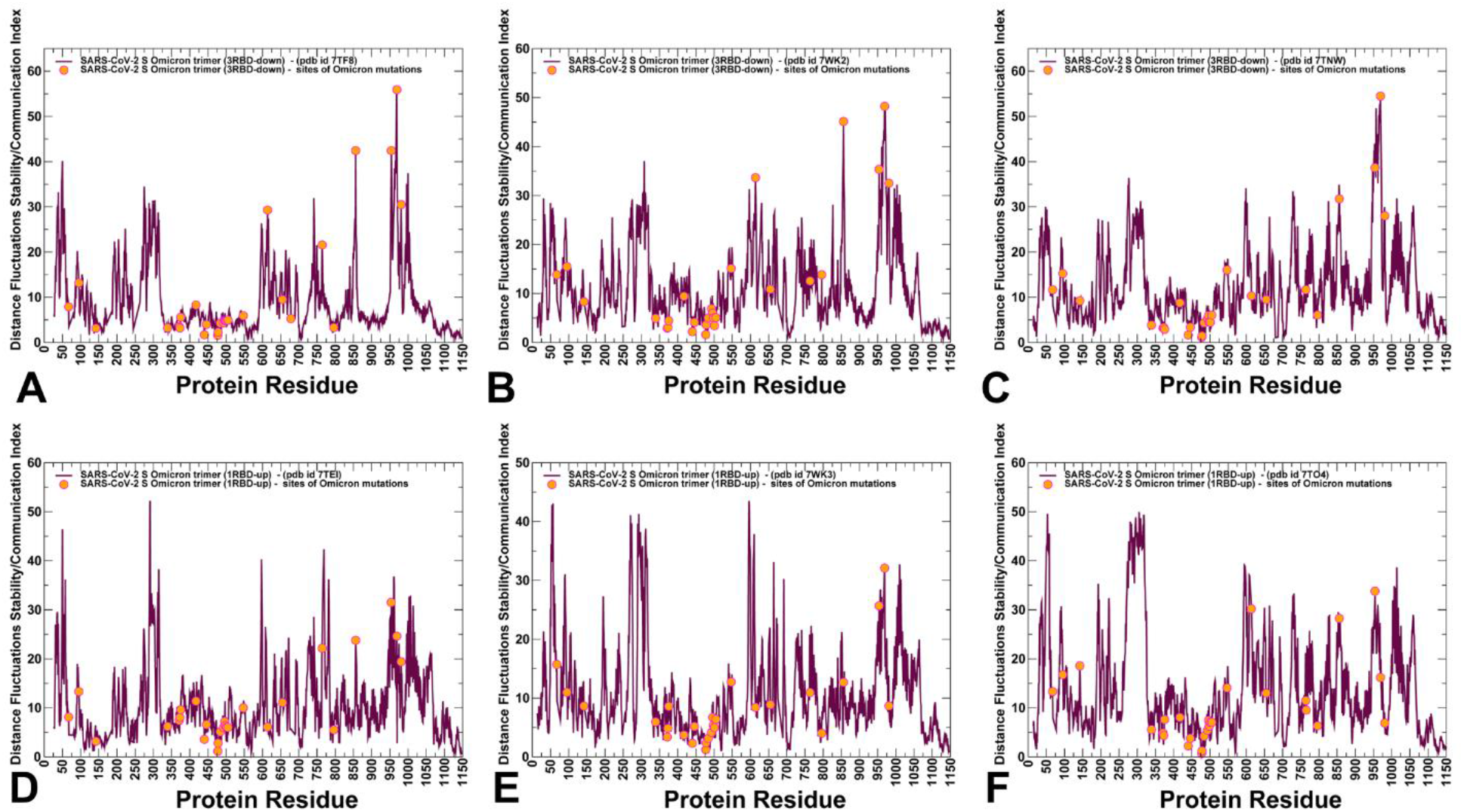
The distance fluctuation stability index profiles obtained from CG-BD simulations of the SARS-CoV-2 S Omicron variant structures. The distance fluctuation stability profiles for the 3RBD-down closed S Omicron structures are shown for pdb id 7TF8 (A), pdb id 7WK2 (B), and pdb id 7TNW(C). The distributions for the 1RBD-up open S Omicron structure are shown for pdb id 7TEI (D), pdb id 7WK2 (E), and pdb id 7TO4 (F). The profiles are shown in maroon-colored lines and the positions of the Omicron mutations A67V, T95I, G142D, G339D, S371L, S373P, S375F, K417N, N440K, G446S, S477N, T478K, E484A, Q493R, G496S, Q498R, N501Y, Y505H, T547K, D614G, H655Y, N679K, P681H, N764K, D796Y, N856K, Q954H, N969K, and L981F are shown in orange-colored filled circles.

**Table 1.** Structures of SARS-CoV-2 S Omicron protein structures examined in this study.

| PDB code | Description | RBDs position |
| --- | --- | --- |
| 7TF8 | SARS-CoV- 2 S Omicron Trimer, closed form | 3 RBD-down |
| 7WK2 | SARS-CoV- 2 S Omicron Trimer, closed form | 3RBD-down |
| 7TNW | SARS-CoV- 2 S Omicron Trimer, closed form | 3 RBD- down |
| 7TEI | SARS-CoV- 2 S Omicron Trimer, open form | 1 RBD-up |
| 7WK3 | SARS-CoV- 2 S Omicron Trimer, open form | 1RBD-up |
| 7TO4 | SARS-CoV- 2 S Omicron Trimer, open form | 1RBD-up |

Using conformational ensembles of the S Omicron structures we computed the fluctuations of the mean distance between each residue and all other protein residues that were converted into distance fluctuation stability indexes that measure the energetics of the residue deformations (Figure 2). The high values of distance fluctuation indexes are typically associated with stable residues as they display small fluctuations in their distances to all other residues, while small values of this parameter would correspond to the flexible sites that experience large deviations of their inter-residue distances. Although the magnitude of the protein residue fluctuations derived from CG-BD simulations may reflect the approximate coarse-grained nature of the energetic force field, the ensemble-averages of these simplified trajectories can yield a robust differentiation of rigid and flexible regions, also pointing to the key dynamic signatures of the Omicron sites. We first analyzed the residue-based distance fluctuation profiles obtained from CG simulations of the S Omicron trimer in the closed state with all 3 RBDs in the down conformation (Figure 2A-C).

Although these profiles displayed some variations in the distribution of highly flexible and stable regions, this analysis revealed several important consistent patterns of structural stability. First, we observed the existence of several dominant and common cluster peaks reflecting topological and dynamical features of the S Omicron trimers in the closed state. The distance fluctuation stability profile for the S1 subunit (residues 14-365) featured a consistent broad corresponding to residues 290-330 in each of the S Omicron closed states (Figure 2A-C). These residues include NTD positions and importantly aligned with the N2R linker (residues 306-334) that connects the NTD and RBD within a single protomer. In the RBD-down conformation a segment of the N2R region forms a β strand (residues 311-318) that interacts with the CTD2 domain (residues 592-685) and β strand (residues 324-328) stacking against CTD1 (residues 528-591).

In the S Omicron closed trimer (Figure 2A), the N2R region secondary structures mediate the intra-protomer salt bridge between R319 and NTD residue E298 and interactions between an important hinge residue F318 against the NTD 295-303 helix. These structural changes are characteristic of both the S Omicron closed trimer (Figure 2A-C) and the open states (Figure 2D-F) resulted in the increased structural stability of the residue cluster 290-320, signaling that this region and N2R linker could play a regulatory role in mediating the long-range dynamics. These observations are consistent with the experimental structural analysis^59^ suggesting that the inter-protomer packing of the RBDs in the and structural rearrangements of the N2R in the RBD-down protomers can define an important mediating element of the stabilizing inter-protomer interactions in the S Omicron closed form. Another significant stability peak in the S1 subunit is broadly aligned with CTD2 region (residues 592-615) that is structurally connected with the N2R linker. These structural stability clusters also highlighted the vital role of anchoring residues F318, F592 and mutational site D614G that remain immobilized in simulations and may be responsible for modulating large movements of the S1 subunit (Figure 2). These results are particularly interesting in the context of our previous all-atom simulation studies of the SARS-CoV-2 S proteins showing that F318 and F592 positions are conserved structurally stable hinge sites of collective motions that can modulate both the inter-protomer and inter-domain changes.^89–94^ According to the distribution profiles of the S Omicron trimers (Figure 2) these positions feature similar dynamic signatures and could play a regulatory role as hinges of global motions.

The important finding of this analysis revealed that the distance fluctuation profiles of the S Omicron closed states displayed the highest stability peaks at the critical residues of the S2 subunit that are aligned with the Omicron mutational positions N764K, N856K, Q954H, N969K, and L981F (Figure 2A-C). The flexible CTD1 region (residues 528-591) is stabilized in both the closed and open S Omicron states via an inter-protomer hydrogen bonding formed by D568 and T572 with the S2 subunit N856K residue. Moreover, the inter-protomer interactions between N969K and Q755 as well as inter-protomer hydrogen bonding of N764K with T315, N317 and Q314 positions in the N2R linker of the adjacent protomer contribute to stabilization of the Omicron mutational sites in the S2 subunit (Figure 2). Hence, strategic position of the S Omicron mutations from S2 subunit can contribute to stabilization of the mobile regions in the S1 subunit and integration of the N2R linker region into the hotspot cluster of structural stability. Hence, these positions emerged as central mechanical hotspots of S protein dynamics that can be related to their biological activity, allosteric control of the inter-protomer interactions and modulation of the binding interfaces. Based on these observations, we suggested that structural stability and long-range interactions in the S Omicron closed trimers may be regulated through a cooperative cross-talk between the Omicron mutational sites in the S2 subunit.

By mapping positions of the Omicron RBD mutations on the distance fluctuation distributions, we observed that the RBD regions are more flexible where N440K, G446S and especially S477N/T478K mutational sites are aligned with the local minima of the distribution (Figure 2A-C). This implies that even in the tightly packed closed trimer the RBD mutational sites maintain a considerable degree of the intrinsic mobility which is required for executing conformational transitions and optimizing RBD interactions with the host receptor. According to the distance fluctuation analysis, conformational transitions from the closed to the open state may be accompanied by the partial redistribution of the stability peaks (Figure 2). Indeed, the relative contribution of the S1 stability clusters becomes more dominant in the S Omicron open states (Figure 2D-F). As a result, in the open S Omicron form the key mechanical sites modulating allosteric interactions would correspond to the inter-domain S1 clusters anchored by F318, F592 and D614 positions. At the same time, the relative contribution of the Omicron mutational sites in the S2 subunit to the distance fluctuation stability profile is reduced, indicating reduced rigidity in these positions which may potentially promote S1/S2 detachment (Figure 2D-F). These results are particularly interesting in the context of the single-molecule biomechanical experiments that quantifies the molecular stiffness of the SARS-CoV-2 S proteins.^128^ According to this study, S protein can exploit mechanical force to enhance its recognition of ACE2 and subsequently accelerate S1/S2 detachment.

To summarize, the results of the dynamics analysis of the SARS-CoV-2 S Omicron structures revealed several important trends. We found that the Omicron mutation positions N764K, N856K, Q954H, N969K, and L981F in the S2 subunit corresponded to important structural stability hotspots in the closed trimers. Based on the analysis, we suggest that these sites could play a regulatory role in modulating balance between protein stability of the closed Omicron trimers and conformational adaptability required to transition to the RBD-up open state. The distance fluctuations stability profiles displayed a partial reduction in rigidity of the S2 Omicron positions in the open states and the concomitant increase of stability in the NTD-RBD inter-domain hinges. Importantly, the Omicron RBD sites are characterized by significant mobility in both closed and open S Omicron trimers, suggesting that the intrinsic conformational plasticity of these RBD positions can be exploited to mediate binding with the host cell receptor. Together, these findings led us to a working hypothesis that the Omicron mutational sites in the S2 subunit and S-RBD regions may form an allosterically coupled interaction network in which structurally stable S2 Omicron sites could act as regulators of the functional movements in the flexible RBD-Omicron sites.

### Conformational and Mutational Frustration in the SARS-CoV-2 S Omicron Structures : Local Frustration of the Omicron Sites Controls Balance of Stability and Plasticity

The functional effects of the Omicron mutations may be characterized by suboptimal solutions to multiple fitness trade-offs that balance protein stability and conformational plasticity to allow for immune escape and binding affinity with the ACE2. To examine how these balancing relationships between structural stability and plasticity are regulated in the S Omicron states, we quantified conformational and mutational frustration of protein residues using the equilibrium conformational distributions (Figure 3). This analysis is based on the ensemble-based profiling of the S Omicron residues by local frustratometer^129, 130^ which computes the local energetic frustration index using the contribution of a residue to the energy in a given conformation as compared to the energies that would be found by mutating residues in the same native location or by creating by changing local conformational environment for the interacting pair.^129–133^ In this approach, a significant stabilization for an individual native pair normalized by the energy fluctuations is considered as indication of minimally frustrated interaction, whereas a destabilizing effect of the native pair deems the corresponding interaction as highly frustrated. The frustration survey of the diverse protein folds showed that minimally frustrated residues may be enriched in the protein core, while the exposed flexible binding interfaces are characterized by the highly frustrated patches that can be alleviated upon binding.^131–133^ Importantly, however, the vast majority of protein residues typically display neutral frustration which implies a moderate degree of stabilization/destabilization for the native interactions as compared to the average energetics induced by conformational or mutational changes in the respective positions.^131–133^

**Figure 3.**
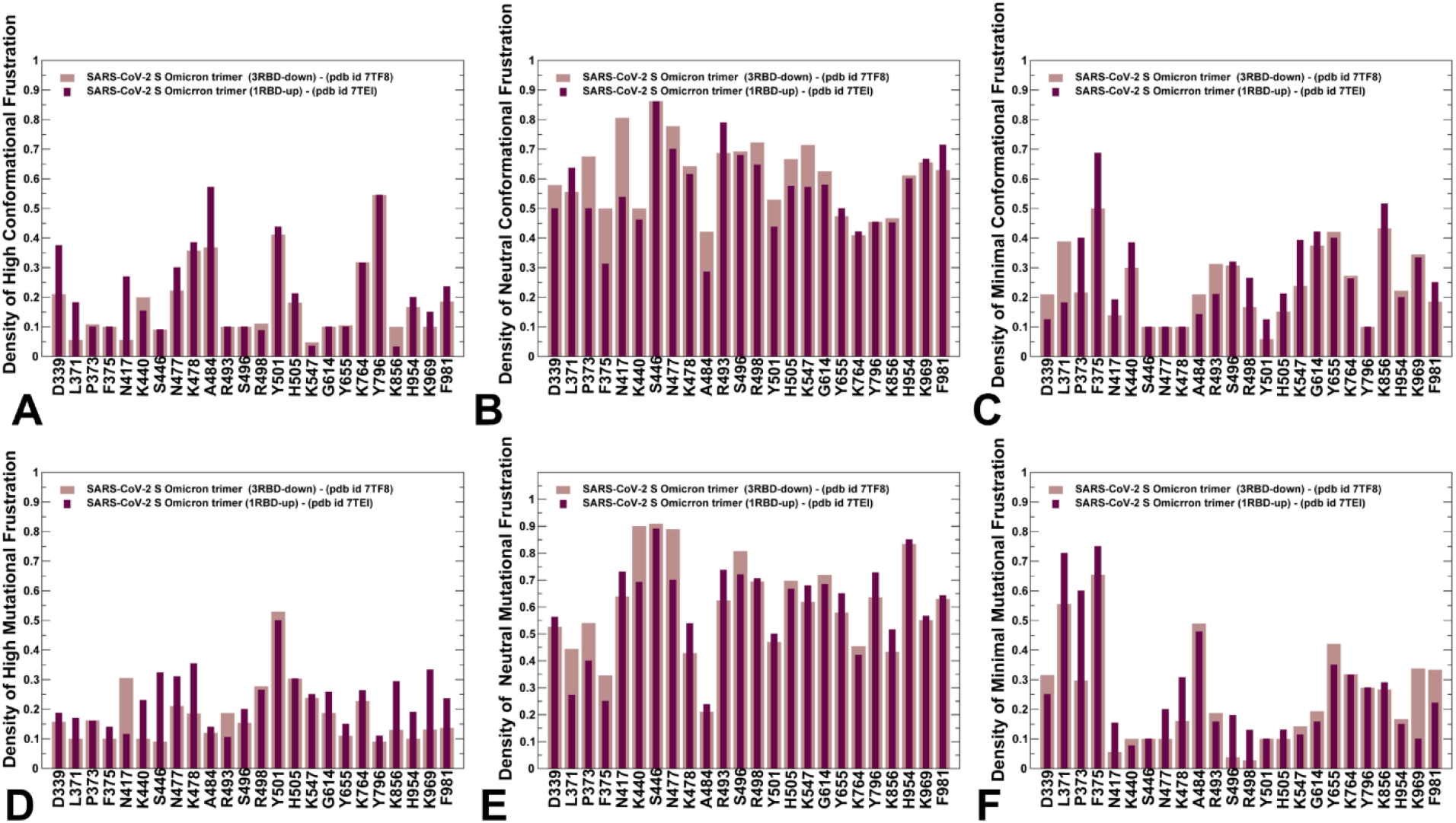
The distributions of conformational and mutational frustration in the SARS-CoV-2 S Omicron trimer structures. (A) The relative densities of high conformational frustration for the Omicron sites in the closed S Omicron structure (brown bars) and open S Omicron structure (maroon bars). (B) The relative densities of neutral conformational frustration for the Omicron sites in the S protein closed and open structures (C) The relative densities of minimal conformational frustration for the Omicron sites in the S protein closed and open structures. (D) The relative densities of high mutational frustration for the Omicron sites in the closed S Omicron structure (brown bars) and open S Omicron structure (maroon bars). (E) The relative densities of neutral mutational frustration for the Omicron sites in the S protein structures. (F) The relative densities of minimal mutational frustration for the Omicron sites in the S protein structures. The distributions were constructed by averaging computations over snapshots from simulations of the S Omicron closed form (pdb id 7TF8, 7WK2, 7TNW) and the S Omicron open states (pdb id 7TEI, 7WK3, 7TO4).

We started by computing the cumulative distribution of the local frustration residue index in the closed and open states by averaging the residue-based local mutational frustration over ensembles of conformations (Supporting Information, Figure S1). The distributions showed the overwhelming preferences for neutral and minimal frustration interactions in the S Omicron structures. The large scale frustration analysis of protein structures previously demonstrated that neutral contacts can be randomly distributed over the protein molecule.^133^ The local frustration analysis of the S Omicron functional states is consistent with a more general frustration analysis of different proteins that revealed the majority of local interactions as neutral (∼50-60%) or minimally frustrated (30%) with only 10% of the total contacts considered as highly frustrated^134^. A similar distribution was found in our recent studies of molecular chaperones, also revealing that conformationally frustrated residues may be involved in local interacting clusters with the less frustrated structurally rigid neighbors.^135^

Despite a different pattern of conformational flexibility in the S1 and S2 subunits of the spike protein, we found that there is a considerable level of neutral frustration contacts in the S Omicron structures which could allow for moderate conformational and mutational variations without compromising the topology and biological activity of the spike protein (Figure 3).

By computing the local density of contacts, we evaluated the distributions of conformational (Figure 3A-C) and mutational frustration indexes for the Omicron mutational sites (Figure 3D-F). The computed values represent the fraction of highly frustrated, neutrally frustrated and minimally frustrated contact density within 5 Å centered in the C_α_ atom from a given residue. The analysis of conformational frustration showed that although the density of highly frustrated contacts is small, several Omicron positions in the RBD region (S477N, T478K, E484A, N501Y, and Y505H) showed an appreciable high frustration density, suggesting that these exposed RBD positions are prone to significant conformational adjustments at the binding interfaces to accommodate the interactions with ACE2 and elicit immune evasion (Figure 3A).

Interestingly, the fraction of high frustration contacts increased for these residues in the open state, reflecting the increased plasticity and accommodating potential of these sites in the receptor-accessible conformations. The noticeable density of highly frustrated contacts was also seen for N764K and D796Y Omicron positions that reside in the exposed regions of S2 subunit (Figure 3A). However, the overwhelming majority of local contacts in the vicinity of Omicron mutational sites are neutrally frustrated, and this pattern is particularly pronounced in the closed S Omicron state (Figure 3B). We also observed appreciable densities of minimally frustrated contacts in the vicinity of the inter-protomer Omicron sites D614G, N856K as well as for S-RBD core positions S371L, S373P, S375F (Figure 3C).

The distribution patterns of mutational frustration for the Omicron sites were generally similar to the conformational frustration indexes (Figure 3D-F). We observed a considerable density of highly frustrated contacts near flexible RBD sites N440K, G446S, S477N, T478K, N501Y, and Y505H (Figure 3D). Interestingly, the high mutational frustration density was especially pronounced for N501Y position, indicating that significant conformational and mutational plasticity in these key sites allow for virus mutability and modulation of binding with ACE2 and antibodies. Similarly, a dominant neutral mutational density was seen for all Omicron sites, particularly for K417N, N440K, G446S (Figure 3E), while a minimal frustration density was significant for the S-RBD core sites S371L, S373P, S375F (Figure 3F).

Collectively, the results of local frustration analysis revealed several important trends reflecting a delicate balance of stability and plasticity in the S Omicron structures. We found that the local density of contacts near the Omicron mutational sites featured a dominant fraction of neutral frustration, while the fractions of highly and minimally frustrated contacts are markedly smaller (Figure 3). This analysis indicated that both S1 and S2 Omicron positions retain an appreciable level of conformational plasticity and mutational adaptability potential, even though S1 domains are characterized by the greater mobility and S2 regions are more rigid in simulations. Since neutrally frustrated contacts can be prevalent in the vicinity of structurally rigid hinge regions, more flexible residues may collocate with the immobilized hinge sites to orchestrate and propagate conformational changes. The observed intrinsic preferences of the Omicron sites for conformational and mutational adaptability may partly explain the emergence of Omicron mutations in the structurally compact S2 subunit. The important implication of the analysis is that a moderate degree of local frustration contacts in the vicinity of the key inter-protomer Omicron positions D614G, N856K, N969K may allow for local repacking and interaction switching (Figure 3). Some of the important inter-protomer contacts stabilized by these sites in the closed S Omicron trimer include N764K-Q314, N856K-D568, N856K-T572, N969K-Q755, and N969K-Y756. According to the local frustration profiles, moderate mutational changes in these sites can be energetically accommodated, leading to redistribution of the inter-protomer bridging pairs that control a balance of stability and plasticity in the S Omicron trimers. Based on these findings, we propose that global allosteric transitions between the closed and open S states may be coordinated through local dynamic changes near sites of Omicron mutations.

To further quantify the contributions of the Omicron sites to the protein stability and examine structural plasticity of the S Omicron trimers, we performed alanine scanning of the inter-protomer interface residues for all studied S protein structures using FoldX approach (Figure 4). The overall distribution density of the free energy changes for the interfacial residues showed a Gaussian-like shape (Figure 4A) where the peak centered around ΔΔG ∼ 0.5 kcal/mol.

**Figure 4.**
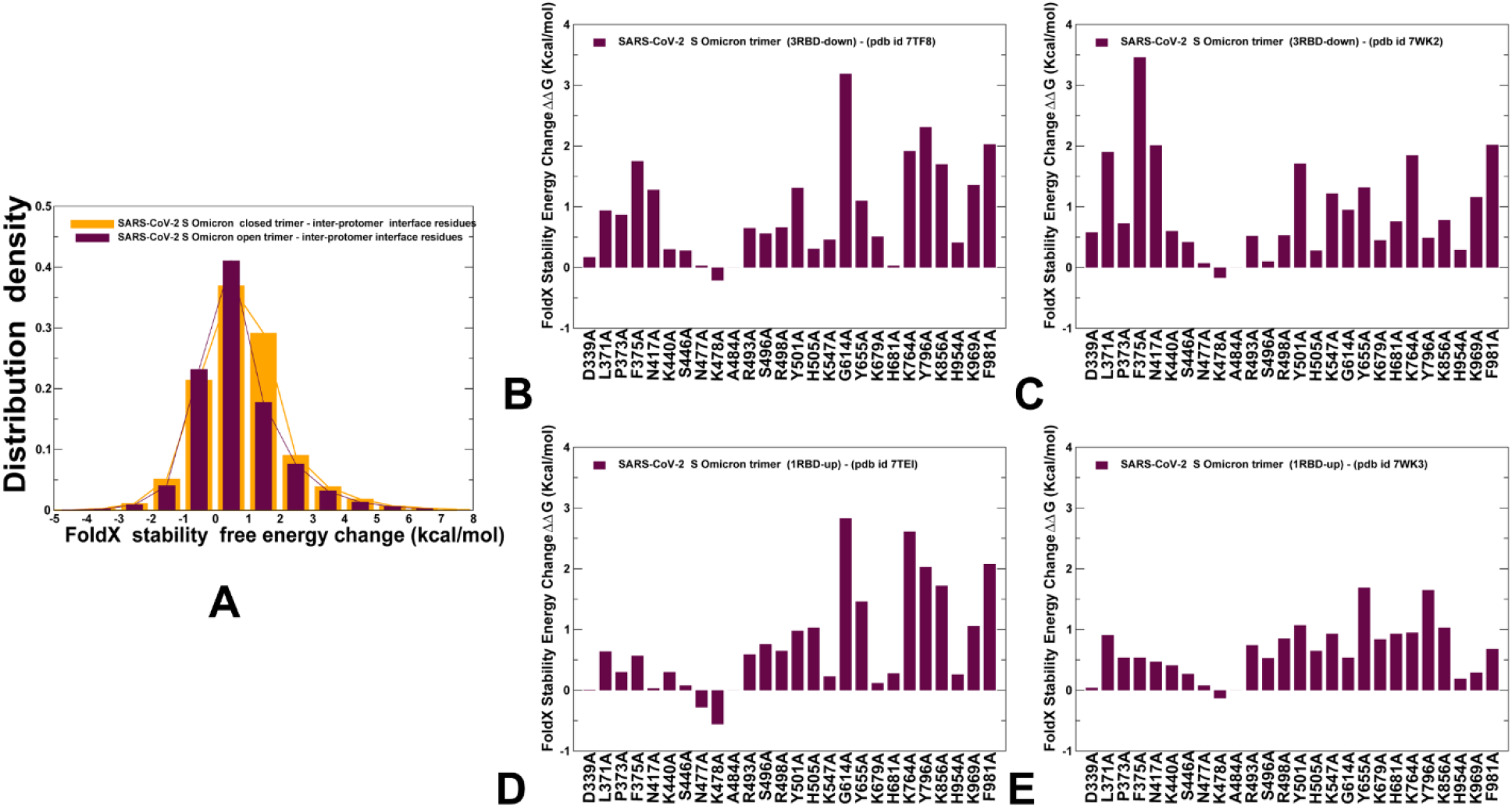
Mutational sensitivity analysis of the inter-protomer residues and alanine scanning of the Omicron sites in the S Omicron closed and open structures. (A) The distribution density of FoldX protein stability changes upon alanine mutations for the inter-protomer contact residues in the S Omicron structures. The stability change density obtained from alanine scanning of the closed S Omicron structures (pdb id 7TF8, 7WK2, 7TNW) is shown in orange bars, and the density obtained from mutational scanning of the 1RBD-up open S Omicron structures (pdb id 7TEI, 7WK3, 7TO4) is shown in maroon bars. The ensemble-based alanine scanning of the Omicron sites in the closed S Omicron structures pdb id 7TF8 (B) and pdb id 7WK2 (C). The ensemble-based alanine scanning of the Omicron sites in the open S Omicron structures pdb id 7TEI (D) and pdb id 7WK3 (E).The protein stability changes are shown in maroon-colored filled bars.

Hence, the vast majority of the free energy changes for the inter-protomer residues are relatively small and destabilizing. This is consistent with the local frustration analysis that employs a completely different force field, suggesting that the inter-protomer contacts in the S Omicron structures are fairly adaptable and tolerant to mutational changes. We also examined the contributions of the Omicron mutational sites to the protein stability (Figure 4B-E). In the closed S Omicron trimers, the largest mutation-induced destabilization changes were observed for the S1 positions S371L, S373P, S375F and S2 positions D614G, N764K, D796Y, N856K, Q954H, N969K, and L981F with ΔΔG ∼ 1.0-1.5 kcal/mol (Figure 4B,C). We also noticed an appreciable sensitivity of the free energy changes to the specifics of the inter-protomer packing in the closed structures. Indeed, larger destabilization changes were seen for the S2 Omicron sites in the tightly packed 3 RBD-down structure (Figure 4B) featuring a dense network of the inter-protomer contacts.^59^

Common to the closed trimer structures, the average destabilization changes for the S1-RBD Omicron positions are small with ΔΔG < 1.0 kcal/mol (Figure 4B,C). Moreover, we found that free energy changes of the highly flexible S477N and T478K sites may be marginally stabilizing, which is consistent with the high frustration of these positions. Despite a more flexible structure of the open trimer, the magnitude of the free energy changes for the Omicron positions was similar to the stability changes in the closed form (Figure 4E,F). These results are also consistent with the local frustration analysis, showing moderate mutational changes and functional plasticity of the Omicron sites, which may be required to modulate multiple fitness trade-offs between stability, binding and immune escape.

### Collective Dynamics of the SARS-CoV-2 S Omicron Structures Reveals Integrating Role of the Omicron Mutational Sites as Major Hinges of Allosteric Conformational Transitions

To identify hinge sites and characterize collective motions in the SARS-CoV-2 S Omicron structures, we performed principal component analysis (PCA) of trajectories derived from CG-BD simulations. The low-frequency ‘soft modes’ are often functionally important as mutations or binding can exploit and modulate protein movements along the pre-existing slow modes to induce allosteric transformations.^136, 137^ The local minima along the slow mode profiles are typically aligned with the hinge centers, while the maxima correspond to the moving regions undergoing concerted movements. Overall, the key functional signature of collective dynamics in the S Omicron structures reflected the preferences for large displacements of the NTD and RBD regions around the inter-protomer hinges (Figure 5). The conserved hinge regions in the closed (Figure 5A-C) and open forms (Figure 5D-F) can regulate the inter-domain movements and functional motions of S1 and S2 regions.

**Figure 5.**
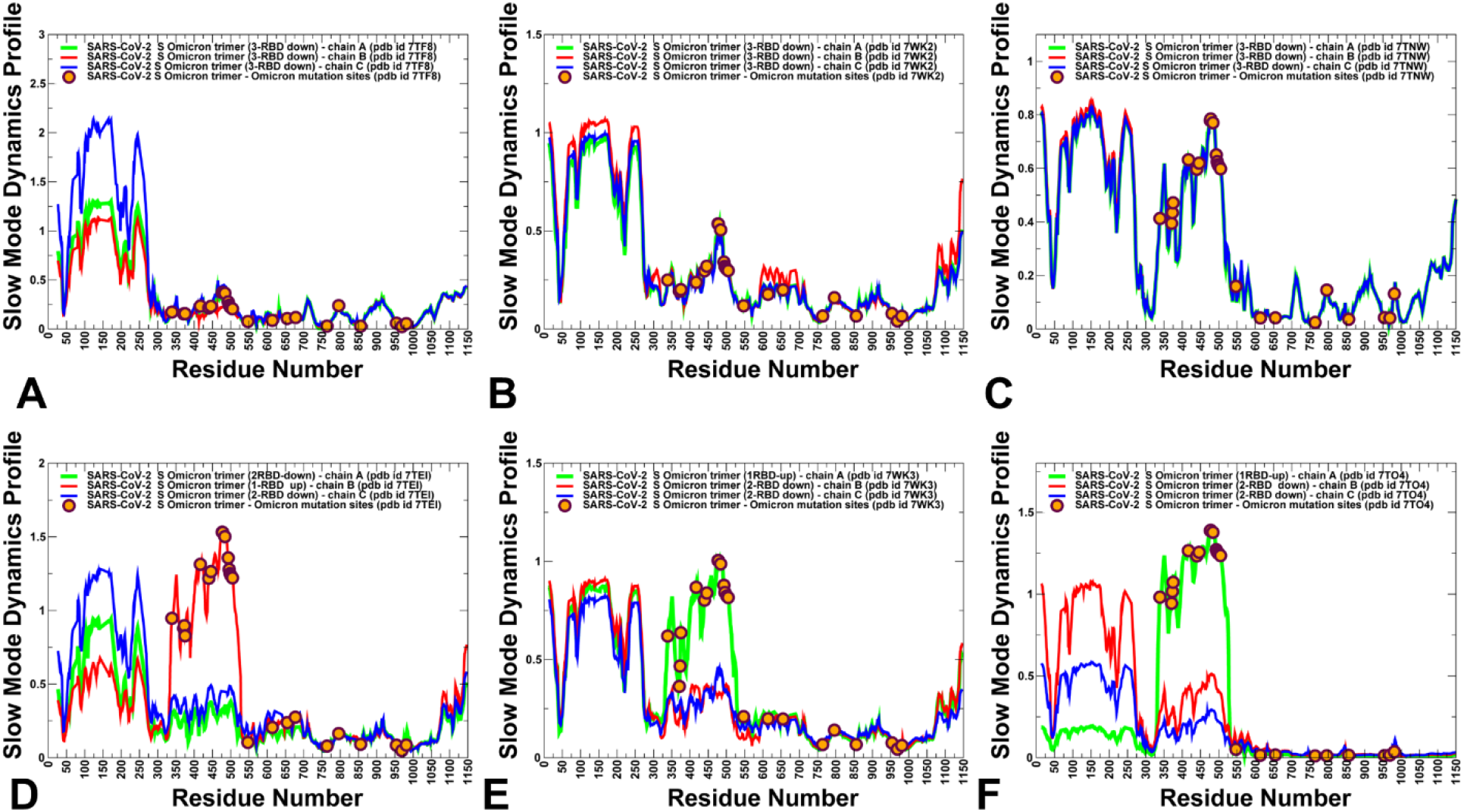
The slow mode mobility profiles of the SARS-CoV-2 S Omicron trimer structures. The slow mode dynamics profiles represent the displacements along slow mode eigenvectors and correspond to the cumulative contribution of the slowest three modes. The slow mode mobility profiles for the closed S Omicron trimer structures pdb id 7TF8 (A), pdb id 7WK2 (B), and pdb id 7TNW (C). The slow mode mobility profiles for the 1RBD-up open S Omicron trimer structures pdb id 7TEI (D), pdb id 7WK3 (E), and pdb id 7TO4 (F). The slow mode profiles for protomer chains A, B and C are shown in green, red and blue lines respectively. The positions of Omicron mutational sites G339D, S371L, S373P, S375F, K417N, N440K, G446S, S477N, T478K, E484A, Q493R, G496S, Q498R, N501Y, Y505H, T547K, D614G, H655Y, N679K, P681H, N764K, D796Y, N856K, Q954H, N969K, and L981F are shown in orange-colored filled circles.

The central and most striking finding of this analysis is a precise alignment of the major hinge positions with the Omicron sites in the S2 subunit (Figure 5). The slow mode profiles in the close S states showed that residues N764K, N856K, Q954H, N969K, and L981F corresponded to the pronounced local minima of the distribution (Figure 5A-C). At the same time, D764Y Omicron position in the S2 subunit is aligned precisely with the local maximum of the profile, indicating that this site participates in collective movements of flexible peripheral regions in the S2 subunit (Figure 5A-C). We further highlighted this observation by zooming the slow mode profiles on the S1-RBD and S2 regions (Supporting Information, Figure S2).

This plot accentuates the major finding showing a clear separation of the Omicron sites into a group of global hinges (N764K, N856K, Q954H, N969K, and L981F), a group of local hinge sites (T547K, D614G, H655Y, N679K) and a group of Omicron residues aligned with the globally moving regions (N440K, G446S, S477N, T478K, E484A, Q493R, G496S, Q498R, N501Y, Y505H) (Figure 5, Supporting Information, Figure S2). The major immobilized hinge positions are conserved and common to both closed and open forms of the S Omicron trimer structure. Several other Omicron sites including T547K, H655Y, and N679K are aligned with local hinge positions, while Omicron RBD sites, and especially RBM positions S477N and T478K corresponded to the pronounced maxima of the distributions indicating their functional role as transmitters of the collective motions (Figure 5). The slow mode mobility profiles in the S Omicron open states highlighted conservation of the key hinge positions in S2 subunit and illustrated the distribution of Omicron mutational sites in the 1 RBD-up protomer (Figure 5D-F). In the open conformations we observed significant functional displacements of the RBD region, indicating that Omicron mutations target sites involved in collective movements. These findings showed that Omicron mutations target almost precisely a specific group of S residues that together determine the functional motions and control global conformational changes. Consistent with our previous studies of the S trimer protein in the native form and other variants^89–92^, the functional dynamics profiles also revealed that F318, A570, I572 and F592 residues are conserved hinge sites that are situated near the inter-domain SD1-S2 interfaces and could function as regulatory centers governing the inter-protomer and inter-domain transitions (Figure 5).

Structural mapping of the Omicron sites on the slow mode dynamics profiles illustrated the key results, showing that the Omicron hinges in the S2 subunit are located along the border between structurally rigid and more flexible compartments of the S protein (Figure 6). The structural map of the collective dynamics in the closed S Omicron trimer highlighted clustering of the Omicron mutational sites into a group of stable regulatory hinges at the inter-protomer interfaces and a group of highly flexible in slow modes S1-RBD positions (Figure 6A). In addition, the remaining Omicron sites in the S2 subunit corresponded to peripheral moving regions. A similar picture is seen in the structural map of the S Omicron open state (Figure 6B), showing consolidation of the Omicron RBD sites in globally moving regions. To further analyze a potential regulatory role of Omicron mutational sites, we characterized the local interaction clusters anchored by the hinge sites F318, A570, T572, F592, D614G, N764K, N856K, Q954H, and N969K (Supporting Information, Figure S3). Instructively, some of these sites participate in direct interaction contacts that can strengthen stability of the hinge clusters. In particular, N856K interacts with T572, while N764K makes favorable contacts T315, N317 and Q314 positions in the N2R linker of the adjacent protomer. N969K is hydrogen-bonded with Y756 and Q755 sites that constitute visible hinge cluster seen in the slow mode mobility profile, while T547K is in direct contacts with S982 and L981F positions.

**Figure 6.**
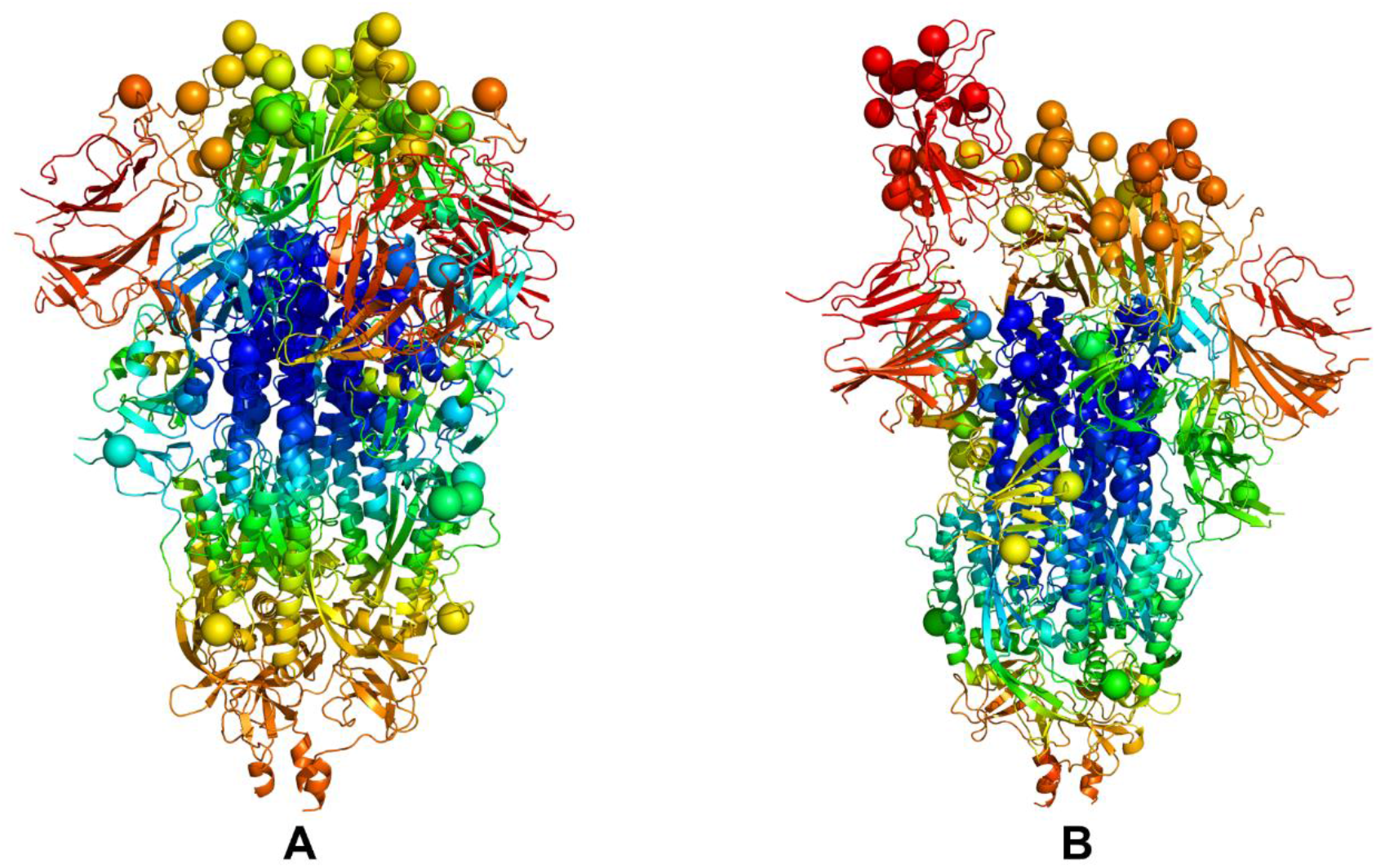
Structural maps of the slow mode mobility profiles for the SARS-CoV-2 S Omicron structures. The structural mapping of the slow mode mobility profiles projected onto the cryo-EM structure of the closed S Omicron trimer, pdb id 7TF8 (A) and the cryo-EM structure of the open S Omicron trimer, pdb id 7TEI (B). The collective dynamics maps are colored according based on the rigidity-flexibility scale with the most rigid regions colored in blue and most flexible regions colored in red. The Omicron mutational sites G339D, S371L, S373P, S375F, K417N, N440K, G446S, S477N, T478K, E484A, Q493R, G496S, Q498R, N501Y, Y505H, T547K, D614G, H655Y, N679K, P681H, N764K, D796Y, N856K, Q954H, N969K, and L981F are shown in spheres for all protomers and colored according to the level of rigidity/flexibility in slow modes.

By mapping these local clusters anchored by Omicron hinge positions (Supporting Information, Figure S3) we observed that these clusters collectively form a tightly packed core at the inter-protomer and inter-domain interfaces (Supporting Information, Figure S3). The Omicron mutations in these regions introduce new stabilizing contacts that promote structural rigidity and expansion of the hinge clusters.

To summarize, the results revealed two groups of Omicron mutational sites that are characterized by distinct and yet highly complementary roles in regulation of allosteric conformational changes that control protein stability and plasticity required for binding and immune response. While a group of Omicron S2 residues may be involved in regulation of global movements and long-range signaling, another group of Omicron sites in the RBD regions can serve as primary receivers of these regulatory signals and execute conformational changes. A clear functional separation of Omicron sites into regulatory hotspots of structural stability that are dominated by S2 residues and allosteric dynamic hotspots in the RBD regions indicated that signaling in the S Omicron structures occurs through coordinated decentralized action in which RBD mutations may differentially affect binding with different partners without sacrificing fidelity of allosteric signaling.

### Dynamic Network Analysis Reveals Role of Omicron Sites as Mediating Bridge Centers of Allosteric Interactions

Our previous studies asserted that SARS-CoV-2 S proteins can operate as functionally adaptable and allosterically regulated machines that exploit the intrinsic plasticity of functional regions to modulate complex dynamic and binding responses to the host receptor and antibodies.^89–94^ Here by using hierarchical network modeling, we characterized the organization of residue interaction networks in the S Omicron functional states. We explore a hypothesis that the Omicron mutations can play an important regulatory role in allosteric interaction networks, where Omicron clusters from S2 and S1 subunits can function as allosterically coupled hotspots that control balance and trade-offs of conformational plasticity and protein stability. We constructed and analyzed the dynamic residue interaction networks using an approach in which the dynamic fluctuations obtained from simulations are mapped onto a graph with nodes representing residues and edges representing weights of the measured dynamic properties. The residue interaction networks in the SARS-CoV-2 spike trimer structures were constructed using a graph-based representation of protein structures in which residue nodes are coupled through both dynamic^117^ and coevolutionary correlations.^118^

We first performed the dynamic network analysis for the S Omicron trimer structures in the closed and open states using the hierarchical community centrality metric (Figure 7). In this network model, allosteric communications are propagated between local communities linked via inter-modular bridging sites. As a result, residues featuring high community centrality values may function as inter-community connectors. The mega-nodes of the higher hierarchical layer aggregate local communities from the lower level where nodes are simply represented by the individual residues. For clarity of presentation and consistency, we report the distributions of community centrality at the lower level where the inter-community bridges are represented by at the single residue level. The high centrality residues are assembled in tight interaction clusters in which the network centrality peaks often align with the hinge regions (residues 315-331, 569-572, 756-760), indicating that these key regulators of functional motions may also mediate communication in the residue interaction networks. The broad centrality distribution reflects the increased number of local stabilizing communities in the closed S Omicron trimer in which many of the RBD mutations stabilize the inter-protomer interfaces and rigidify the 3-RBD-down form. A broad distribution of the mediating regions with medium-to-high centrality implies that S1 and S2 subunits in the S Omicron closed states may be linked through a wide ensemble of distinct communication routes, thus indicating robustness of signal transmission. Importantly, we found that for all protomers in the closed state, the dominant centrality peak corresponds to the N2R linker region (residues 300-336) that connects the NTD and RBD within a single protomer unit. CTD1 region and strong centrality density in the NTD regions positions (Figure 7A,B).

**Figure 7.**
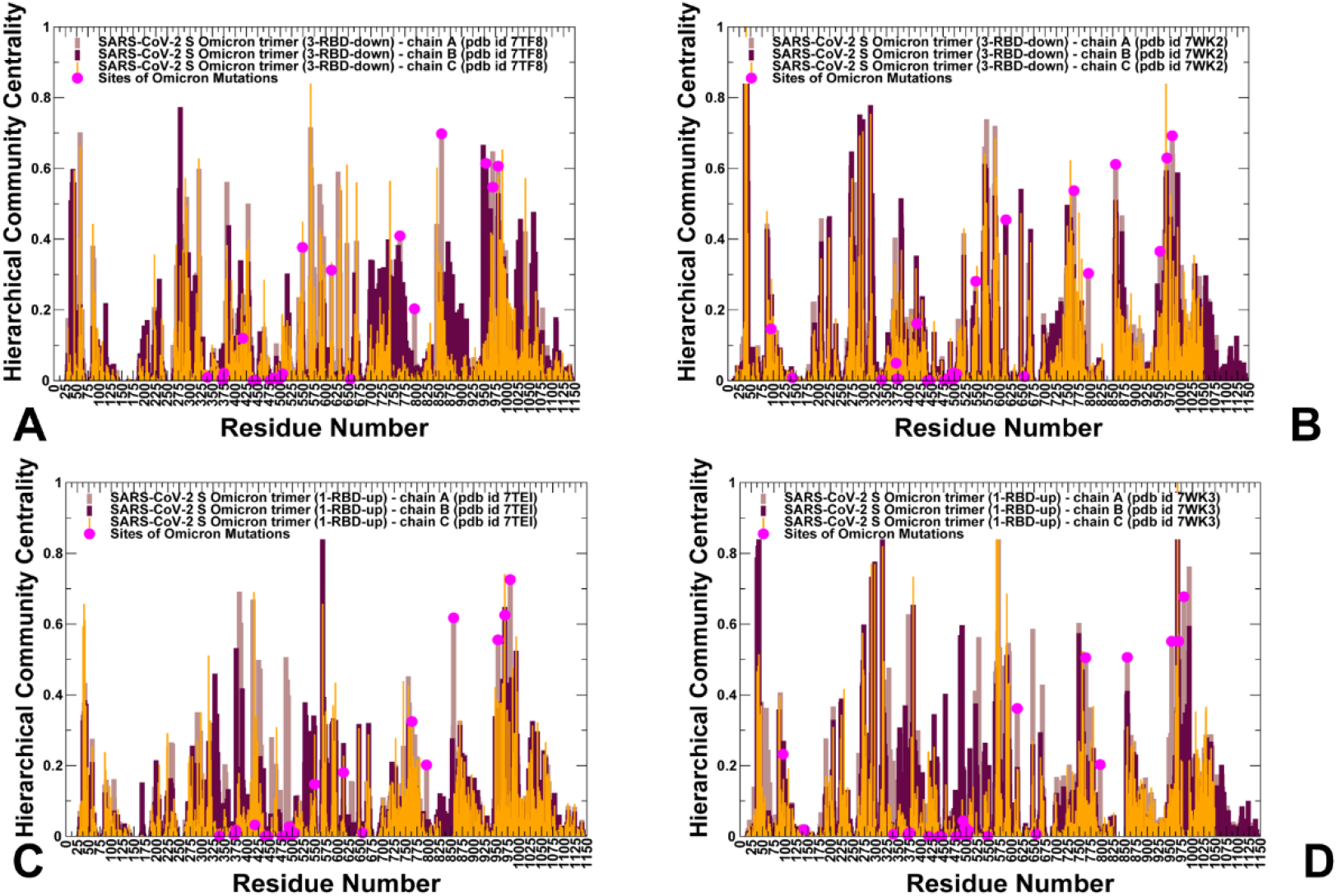
Hierarchical community centrality profiles for the SARS-CoV-2 S Omicron structures The community centrality profiles for the closed S Omicron trimer structures, pdb id 7TF8 (A) and pdb id 7WK2 (B). The community centrality profiles for the open S Omicron trimer structures, pdb id 7TEI (C) and pdb id 7WK3 (D). The distributions are shown for protomer A in brown bars, for protomer B in maroon bars, and for protomer C in orange-colored bars. The positions of the Omicron sites G339D, S371L, S373P, S375F, K417N, N440K, G446S, S477N, T478K, E484A, Q493R, G496S, Q498R, N501Y, Y505H, T547K, D614G, H655Y, N679K, P681H, N764K, D796Y, N856K, Q954H, N969K, and L981F are shown in magenta-colored filled circles.

By mapping the positions of the Omicron mutational sites onto the network profiles, we observed that the Omicron S1-RBD sites featured low centrality values, whereas S2 Omicron positions T547K, N764K, N856K, N969K and L981F exhibited high centrality and are aligned with the maxima along the distribution (Figure 7A,B). These findings suggested that these two clusters of Omicron mutations could play distinct and yet complementary roles in allosteric communications in the S trimer structures. Indeed, the high community centrality for key Omicron positions at the inter-protomer hinge regions supports our conjecture that these sites play role of regulatory allosteric centers and stability hotspots in the S protein that control signaling in the closed the S protein. The low-to-medium centralities and high flexibility of the S1-RBD Omicron sites pointed to their role in modulating conformational rearrangements and functional adaptability of the RBD regions.

The important finding of the centrality analysis is the fact that the maxima of the distribution profile are largely associated with the residues involved in the inter-protomer bridges N764K-Q314, S982-T547K, N856K-D568, N856K-T572, N969K-Q755, N969K-Y756, S383-D985, F855-G614, V963-A570, N317-D737, R319-D737, R319-T739, R319-D745, and K386-F981 (Figure 7A,B and Figure 8). Many of these mediating inter-protomer bridges are anchored by the Omicron mutational sites (T547K, D614G, N764K, N856K, N969K, and L981F) from the S2 subunit that are also associated with hinge regions as evident from the collective dynamics analysis. The inter-protomer network of contacts between the RBDs is formed by low-to-medium centrality residues that are mobile and allow for structural rearrangements (Figure 8). The moderate centrality of the RBD residues involved in the inter-protomer interactions implies that fidelity of allosteric signaling in the S closed trimer can be tolerant to conformational changes and network rewiring in these regions.

**Figure 8.**
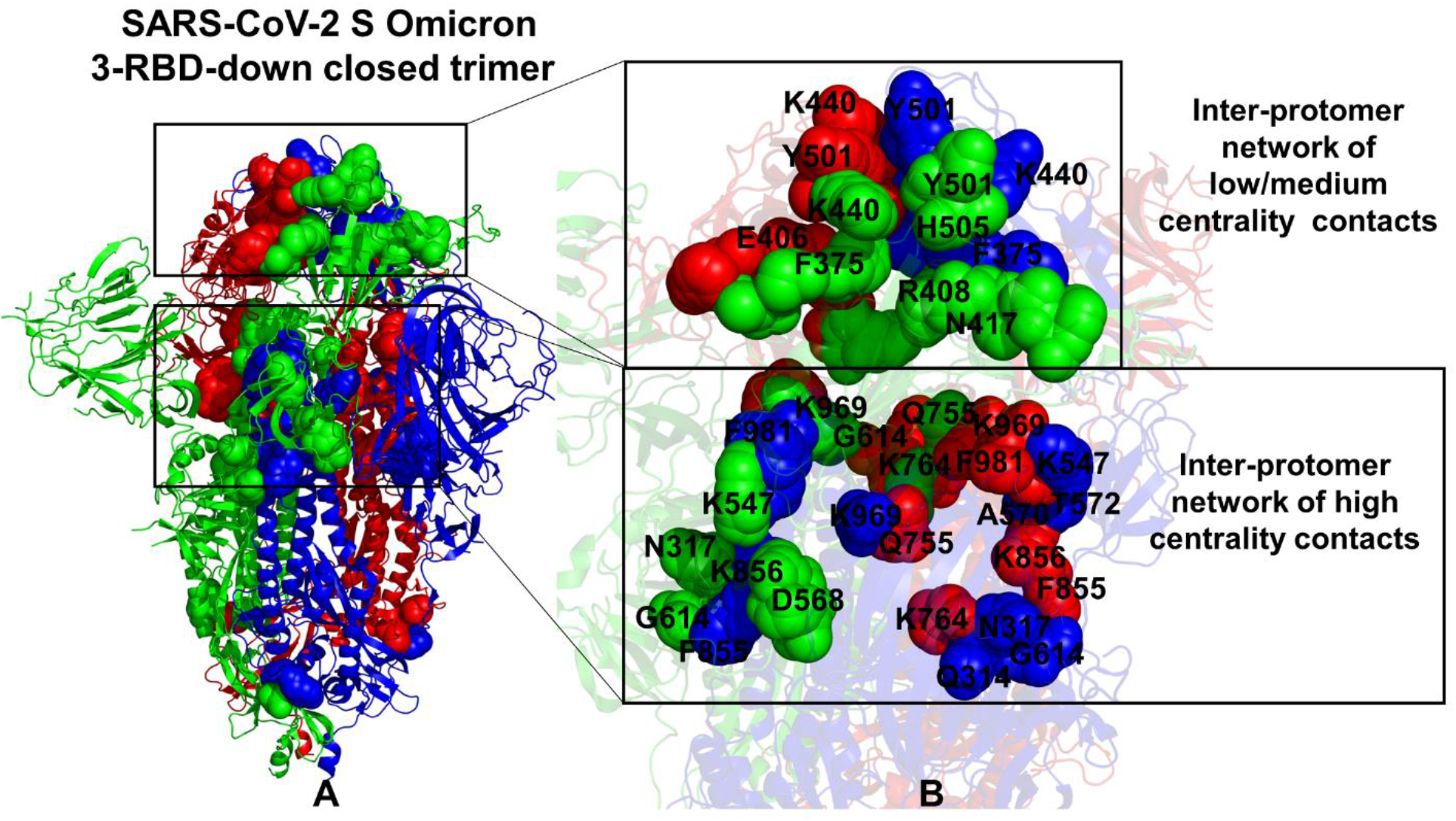
Structural mapping of the inter-protomer network bridges in the 3-RBD down closed S Omicron trimer structures (A) The overall view of the major inter-protomer network bridges. The closed S Omicron trimer is shown in ribbons with the protomers A,B,C in green, red and blue respectively. The inter-protomer bridges are shown in spheres. (B) A close up of the inter-protomer RBD-RBD network is formed by low centrality residues. (top of panel B). The inter-protomer network of high centrality bridging sites (bottom of panel B). The residues that form the inter-protomer network are shown in spheres colored according to the respective protomer and annotated.

These results provide an additional insight to structural studies^56, 59^ showing how Omicron substitutions in SD1 and S2 may enhance the allosteric network between the neighboring protomers that stabilize the closed Omicron structure. Recent structural studies^59^ raised the important question concerning the trade-offs in the S Omicron protein between a potential for conformational changes and the increased stability of the closed state. Our network analysis suggested that the inter-protomer stabilization of high centrality contacts is the main determinant of the S protein stability, while the inter-protomer contacts in the RBD regions are more conformationally adaptive and less network-sensitive.

The residue centrality profile for the 1RBD-up open state featured only a partial redistribution of several mediating clusters, revealing the dominant centrality peak aligned with the N2R linker region (residues 300-336) for the down protomer B (Figure 7C,D). The overall shape of the centrality distribution is preserved in the open S Omicron forms, in which the local peaks are aligned with the residues involved in the inter-protomer interactions. The inter-protomer bridges include N764K-Q314, S982-T547K, N856K-D568, N856K-T572, N969K-Q755, N969K-Y756, N317-D737, R319-D737, R319-D745, R319-T739 are preserved in the interaction network of the open state (Figure 9). In general, our analysis showed that structural and dynamic changes between the closed and open S Omicron states may not induce drastic rearrangements in the interaction networks but rather modulate the strength of the inter-protomer bridging centers. Interestingly, in the 1RBD-up protomer, the repulsive inter-protomer contacts of the closed states K378-R408 and K386-F981 are replaced by the inter-protomer pairs K378-Q493, K386-Y421, Y489-L371, and Y369-N417 that strengthen RBD-RBD contacts of the down protomers (Figure 9).

**Figure 9.**
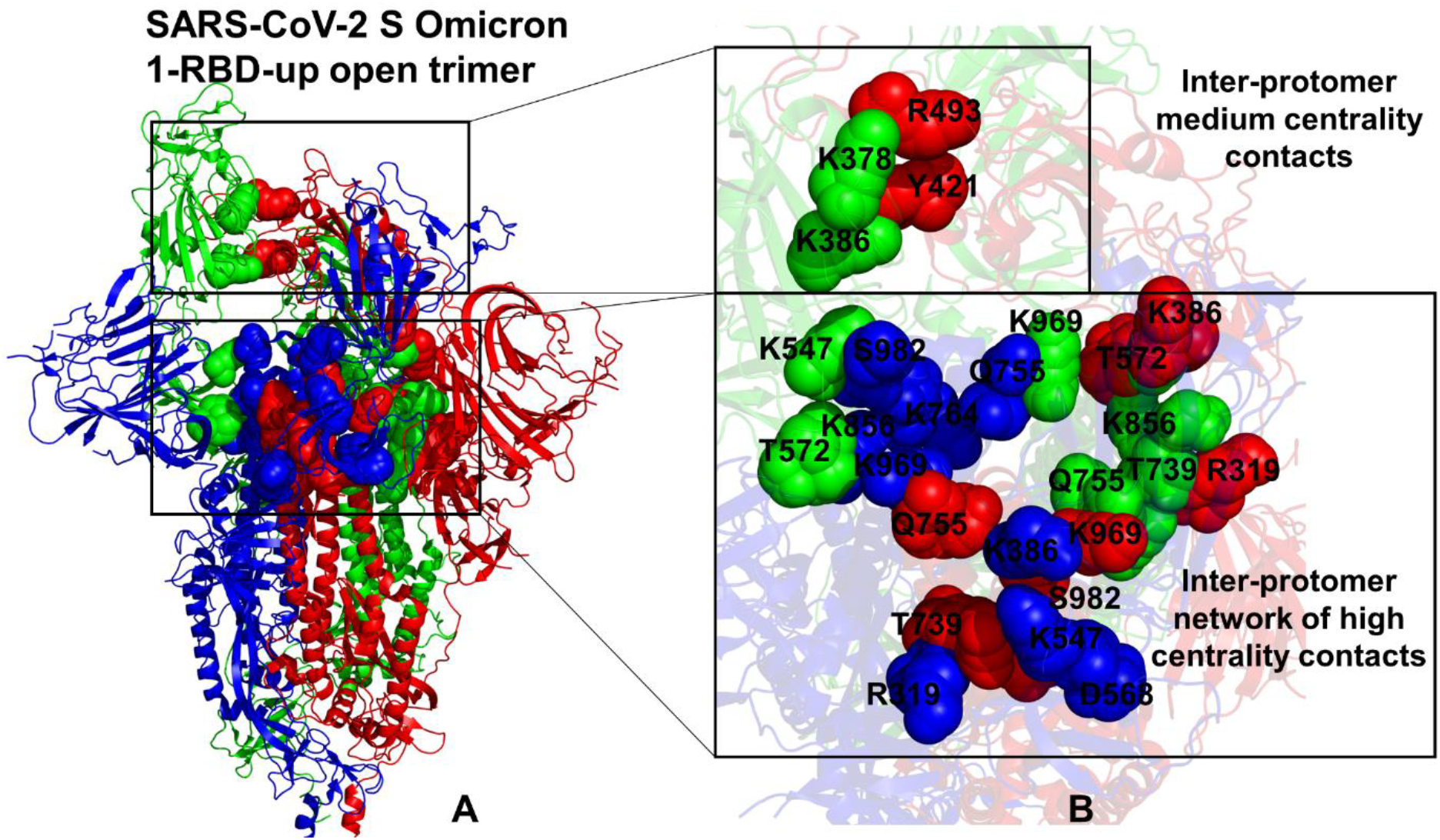
Structural mapping of the inter-protomer network bridges in the 1RBD-up open S Omicron trimer structures (A) The overall view of the major inter-protomer network bridges. The open S Omicron trimer is shown in ribbons with the protomers A,B,C in green, red and blue respectively. The inter-protomer bridges are shown in spheres. (B) A close up of the inter-protomer RBD contacts is formed by low centrality residues (top of panel B). The inter-protomer network of high centrality bridging sites(bottom of panel B). The residues that form the inter-protomer network are shown in spheres colored according to the respective protomer and annotated.

The result of network analysis also highlighted an important role of electrostatic interactions mediated by the Omicron mutations that can modulate the strength of the inter-protomer couplings. The cumulative effect of the electrostatic bridges formed by lysine mutations in the Omicron positions T547K, N856K, N764K, and N969K results not only in the enhanced direct inter-protomer contacts but also through stabilization of the global allosteric network. These findings are consistent with recent studies showing that the breakage of several RBD–S2 electrostatic interactions is required for S1–S2 dissociation.^137^ Our observations are also in accord with the more rigorous quantum chemical interaction analyses of SARS-CoV-2 S protein based on fragment molecular orbital calculations.^138^ Consistent with our findings, this study demonstrated a crucial role of charged Lys residues in differentiating the inter-protomer interactions of the closed and open S protein forms.^139, 140^ Another insightful atomistic simulation study of the kinetics of conformational changes in the SARS-CoV-2 and CoV-1 S proteins showed that a salt bridge R319-D745 between the RBD of the active up protomer and the S2 region in the CoV-2 S protein contributes to the relative stability of the active spike state and the differential dynamic behavior between SARS CoV-2 and SARS CoV-1 spike trimers.^141^

To summarize, the network modeling demonstrated that the Omicron mutations from the S2 subunit serve as mediating network hotspots that form stable inter-protomer electrostatic bridges connecting local stable communities and therefore could function as allosteric switches of signal transmission. This analysis also indicated that the inter-protomer contacts in the RBD regions of the closed trimer are more conformationally adaptive, where the low centrality of the S1-RBD Omicron sites allow for network rearrangements between the closed and open forms.

### Modeling of Allosteric Communication Paths and Diffusion on the Residue Interaction Network using Markov Transient Analysis

We suggest that the inter-protomer interaction pairs featured as key bridges of allosteric interactions may operate as a network of regulatory switches that could cooperatively control signal transmission in the spike protein. The existence of diverse inter-modular network bridges at the inter-protomer regions may also indicate that allosteric signaling can be transmitted through an ensemble of communication pathways rather than through a unique allosteric route connecting S2 regions with the RBD.

By leveraging the residue interaction graph representation of the network, we utilized Markov transient analysis of propagation of a random walker on the network nodes to model allosteric pathways and examine the kinetics of diffusion process on the residue interaction graph by initiating process from the key inter-protomer hinge centers (Figure 10). For simulations, we selected as sources of random diffusion on the protein network the hinge residues A570 and T572 that were shown to play a critical role in allosteric regulation of the spike proteins in both computational studies^89–94^ as well as in structural and biophysical experiments.^39^ Using the evolution of a random walker on the interaction graph and Markov analysis, we obtained the distributions of the transient path times as a function of the distance from the selected communication source (Figure 10). Through this analysis we examine the network connectivity and communication paths in the residue interaction network as well as the distribution of rapidly reachable protein residues that can serve as a communication map of the regulatory signals. The profiles revealed several interesting trends in allosteric communication paths and important differences between allosteric connections in the closed and open S Omicron states. In the closed form, we observed a more “compressed” distribution shape showing that most of the S residues can be efficiently reached from the major inter-domain hinge (Figure 10A). There is a noticeably smaller dispersion in the transient times at a long distance (30Å −60 Å) for the closed trimer (Figure 10A) as compared to the 1RBD-up open form in which the longer transient times are observed (Figure 10B). This is consistent with and the network centrality analysis showing a broad distribution of communication paths in the closed S Omicron trimer (Figure 10A,C). By highlighting the transient times data points for the Omicron sites, it could be seen that the signal can rapidly propagate between the inter-protomer hinges and positions of the Omicron mutations. The distribution of transient times to Omicron sites aligned with the most rapidly communicating residues in the system (Figure 10A). Noticeably, even the transient times required to reach the remote RBM Omicron sites residing at distances > 50 Å from the source are shorter that for other remote distant spike residues. A generally similar trend of rapid communication between the regulatory inter-protomer site and Omicron positions was observed in the open trimer (Figure 10B). In this case, we also found that the Omicron positions can be rapidly reached from the regulatory hinges. Structural mapping of rapidly communicating sites in the 1 RBD-up open form displayed a smaller and more localized network of paths between the inter-protomer hinge source and S1 regions (Figure 10D) where the 1 RBD-up domain may be connected to the source through an efficient communication route. The topography of these maps showed significant differences in the distribution and density of major communication routes connecting S1 and S2 regions in the functional states of the S trimer. The communication network of rapidly reachable residues in the closed state featured a dense network with many alternative global paths that engage the inter-protomer bridges mediated by the Omicron sites (Figure 10A, C). The connectivity of communication hubs is high in the closed form reflecting the broader allosteric network that may enable efficient and robust signal transmission. The number of hubs is reduced in the open S trimer making allosteric signaling more dependent on several key inter-protomer bridges governing communications. We argue that the dense allosteric network of high centrality sites and communication hubs seen in the closed form could allow for diverse alternative routes between S2 and S1 regions, thereby protecting the robustness of allosteric signaling. At the same time, allosteric communications in the open form may become more sensitive and vulnerable to targeted mutations of the key mediating centers.

**Figure 10.**
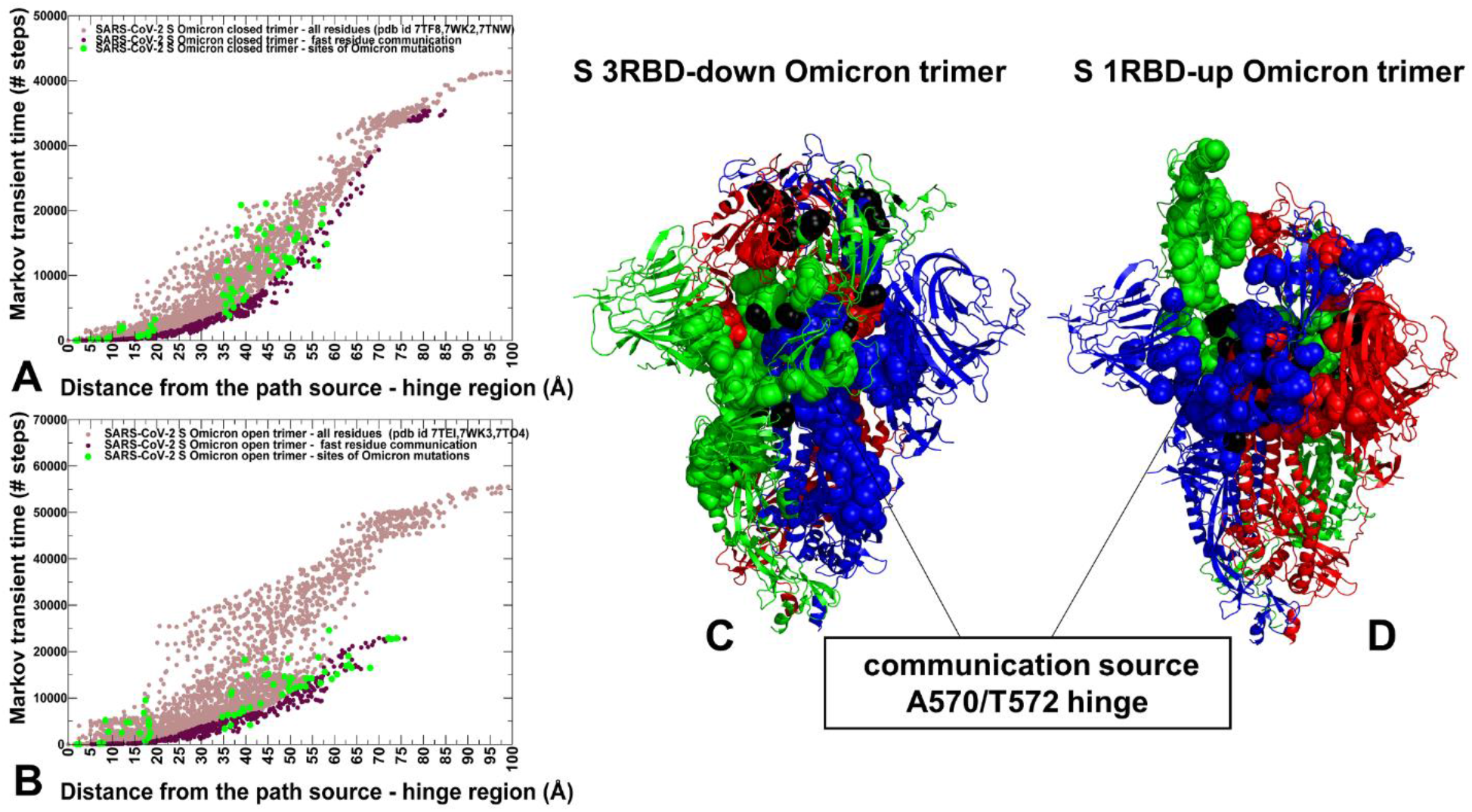
Analysis of allosteric communications in the SARS-CoV-2 S Omicron trimers. The distributions of the Markov transient times for the paths originated from the source hinge region A570/T572 in the protomer A and reaching other trimer residues as a function of the distance from the source are shown for the S Omicron closed trimer (A) and S Omicron open trimer (B). The data points for the average transient times for protein residues are shown in light brown circles. The rapidly communicating sites with the fastest transient times from the source hinge region are shown in maroon-colored circles. The transient times data points corresponding to the Omicron mutation sites in all three protomers are shown in green-colored circles. Structural mapping of the major fast communicating sites with the source hinge region A570/T572 are shown for the closed trimer (C) and open trimer (D). The S Omicron trimer structures are shown in ribbons, with the protomer A in green, protomer B in red, and protomer C in blue. The fast communicating trimer residues are shown in spheres colored according to the respective protomer. The Omicron mutational sites on these structural maps are shown in black-colored spheres. The position of the path communication origin which is selected for these simulations and corresponds to the major hinge region A570/T572 is indicated by the arrows and annotated.

To summarize, the central finding of this analysis is the emergence of efficient signal communication between the regulatory regions driving functional spike transitions and sites of Omicron mutations. The results showed that Omicron mutations at the key regulatory positions can determine topography of signal communication pathways operating through state-specific cascades of control switch points. Hence, the Omicron sites may be involved in a cross-talk with the central regulatory hotspots and enable allosteric control and modulation of structural stability and conformational changes which is central for spike activation and virus transmissibility.

## Conclusions

In this work, we combined molecular simulations of multiple full-length SARS-CoV-2 S Omicron trimers structures in the closed and open states with the local frustration analysis of conformational ensembles and dynamic modeling of allosteric interaction networks to understand specific functional roles of the Omicron mutations in regulating the balance and trade-offs between protein stability and conformational adaptability. We found that the Omicron mutation positions N764K, N856K, Q954H, N969K, and L981F in the S2 subunit are important structural stability hotspots in the closed trimers that play a regulatory role in spike activation. Conformational and mutational frustration profiling of the S Omicron structures showed the dominant presence of moderately frustrated contacts at the inter-protomer regions that enables a level of energetic tolerance to local conformational changes near the regulatory Omicron positions and provides the energetic driver for allosteric transitions between the closed and open states. The ensemble-based alanine scanning of the Omicron positions supported the local frustration analysis, showing moderate destabilization changes at the inter-protomer Omicron sites which may be required to modulate the trade-offs between stability, binding and immune escape. The important finding of this study is that Omicron sites can be separated into a group of structural stability hotspots from the S2 subunit and locally frustrated dynamic centers in the S1-RBD regions that mediate large conformational changes of flexible regions. Through an extensive network modeling, we found that the inter-protomer bridges anchored by the S2 Omicron sites connect local stabilizing communities, showing that Omicron positions are important for efficiency of the global allosteric network. We further examined the nature of allosteric communications and diffusion process on the residue interaction graph using Markov transient analysis. The important finding of this analysis is that regulatory hinges and the Omicron positions are efficiently inter-connected by fastest allosteric pathways and can cooperate to adaptively modulate the regulatory functions and activation of the S protein. The results of this study are consistent with structural and biophysical experiments, revealing specific roles of the Omicron mutational sites that enable allosteric modulation of structural stability and conformational changes which are central for spike activation and virus transmissibility. The insights from this investigation suggest therapeutic venues for targeted exploitation of the binding hotspots and allosteric communication centers that may potentially aid in evading drug resistance.

## SUPPORTING INFORMATION

Figure S1 describes the distribution of the conformational and mutational frustration in the closed and open S Omicron protein states. Figure S2 presents the slow mode profiles of the S Omicron conformational dynamics highlighting the distributions in the S1-RBD and S2 regions. Figure S3 depicts structural maps of the local clusters anchored by the Omicron hinge sites at the inter-protomer interfaces. This material is available free of charge via the Internet at http://pubs.acs.org.

## AUTHOR INFORMATION

* **Corresponding Author**

Phone: 714-516-4586; Fax: 714-532-6048; The authors declare no competing financial interest.

## Acknowledgment

This work was partly supported by institutional funding from Chapman University. The author acknowledges support by the Kay Family Foundation Grant A20-0032.

## ABBREVIATIONS

SARS: Severe Acute Respiratory Syndrome
RBD: Receptor Binding Domain
ACE2: Angiotensin-Converting Enzyme 2 (ACE2)
NTD: N-terminal domain
RBD: receptor-binding domain
CTD1: C-terminal domain 1
CTD2: C-terminal domain 2.

