## Supporting Figures S1-S3 for "Hierarchical Computational Modeling and Dynamic Network Analysis of Allosteric Regulation in the SARS-CoV-2 Spike Omicron Trimer Structures: Omicron Mutations Cooperate to Allosterically Control Balance of Protein Stability and Conformational Adaptability"

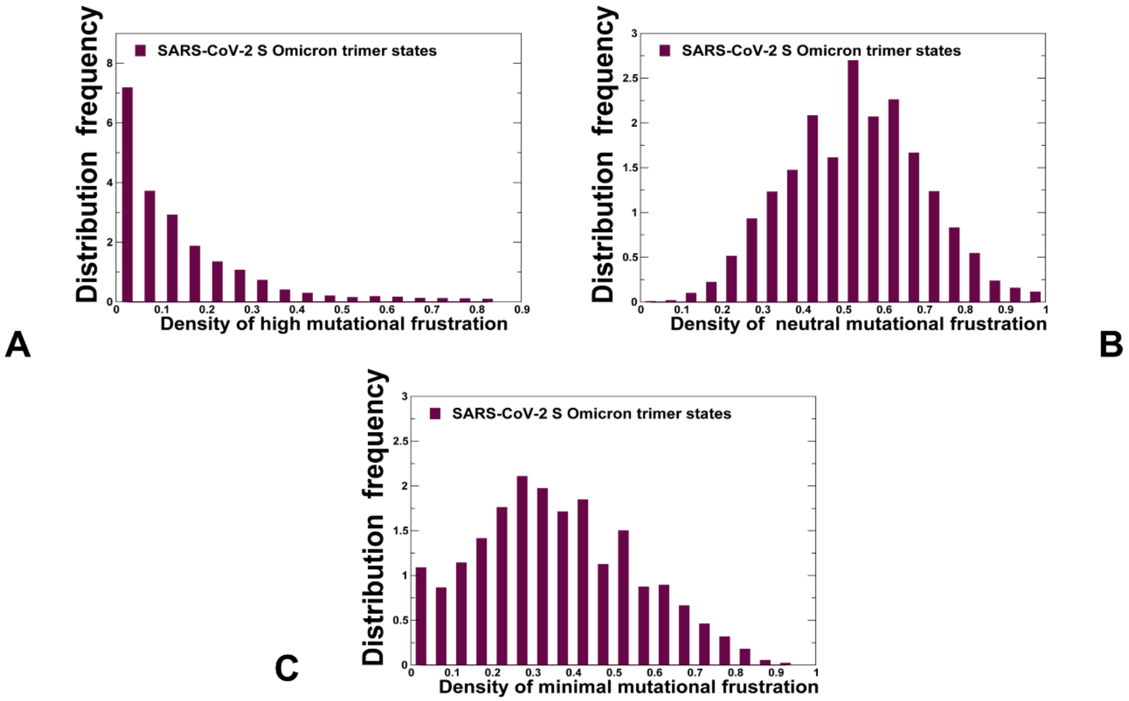


**Figure S1.** The distribution of local mutational frustration in the closed and open forms of the SARS-CoV-2 S Omicron structures. The relative density of highly frustrated contacts (A), neutrally frustrated contracts (B) and minimally frustrated contacts (C) in the S Omicron closed and open trimer structures. The distributions were constructed by averaging computations for the closed S Omicron trimer states (pdb ids 7TF8, 7WK2, and 7TNW) and open S Omicron trimers (pdb ids 7TEI, 7WK3, and 7TO4).

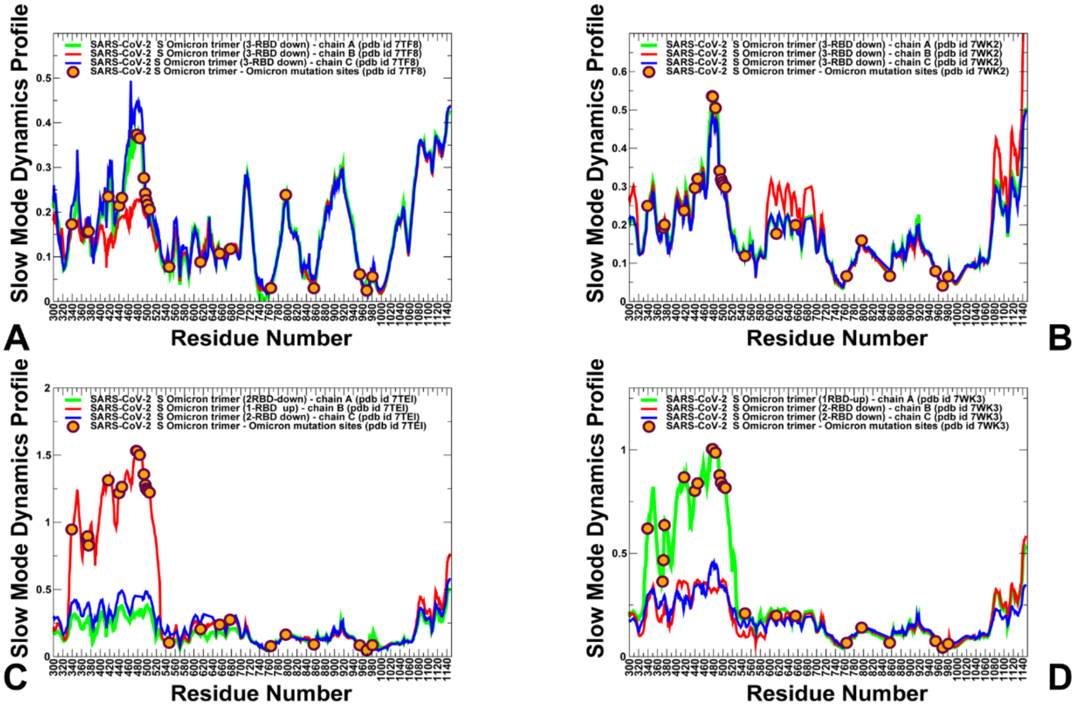


**Figure S2.** A close-up of collective dynamics profiles for the SARS-CoV-2 S Omicron trimer structures highlights the distributions in the S1-RBD and S2 regions The slow mode dynamics profiles represent the displacements along slow mode eigenvectors and correspond to the cumulative contribution of the slowest three modes. The slow mode mobility profiles for the closed S Omicron trimer structures pdb id 7TF8 (A), pdb id 7WK2 (B). The slow mode mobility profiles for the 1RBD-up open S Omicron trimer structures pdb id 7TEI (C), pdb id 7WK3 (D). The slow mode profiles for protomer chains A, B and C are shown in green, red and blue lines respectively. The positions of Omicron mutational sites G339D, S371L, S373P, S375F, K417N, N440K, G446S, S477N, T478K, E484A, Q493R, G496S, Q498R, N501Y, Y505H, T547K, D614G, H655Y, N679K, P681H, N764K, D796Y, N856K, Q954H, N969K, and L981F are shown in orange-colored filled circles.

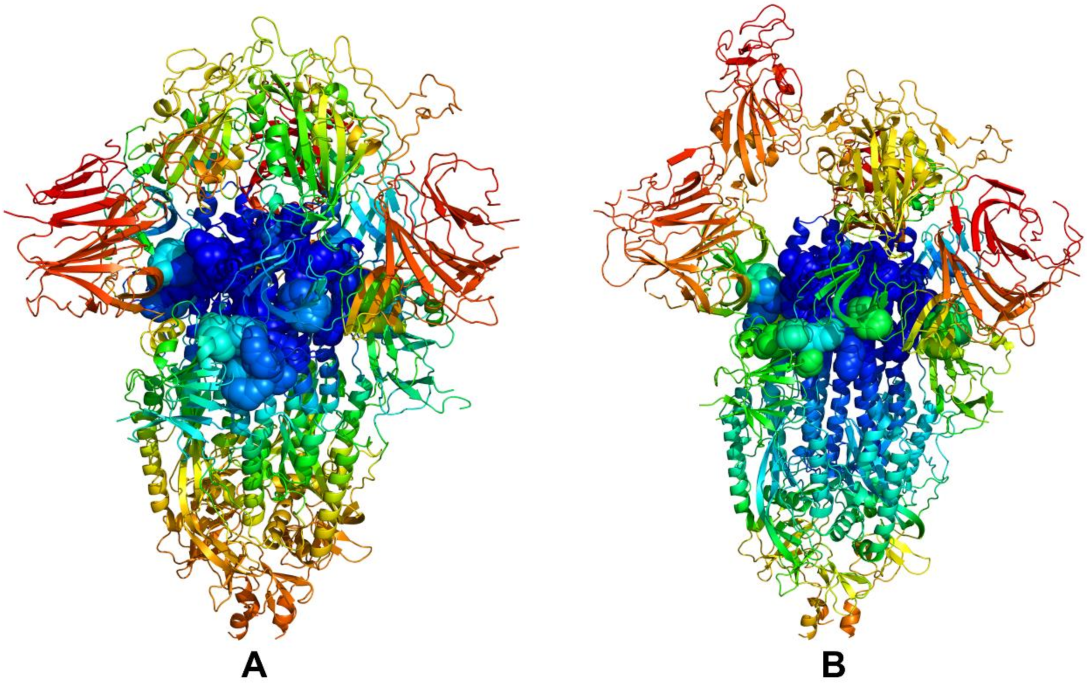


**Figure S3.** Structural mapping of local interaction clusters anchored by the hinge sites F318, A570, T572, F592, D614G, N764K, N856K, Q954H, and N969K. The structural mapping of the slow mode mobility profiles projected onto the cryo-EM structure of the closed S Omicron trimer (pdb id 7TF8) (A) and the cryo-EM structure of the open S Omicron trimer (pdb id 7TEI). The collective dynamics maps are colored according based on the rigidity-flexibility scale with the most rigid regions colored in blue and most flexible regions colored in red.

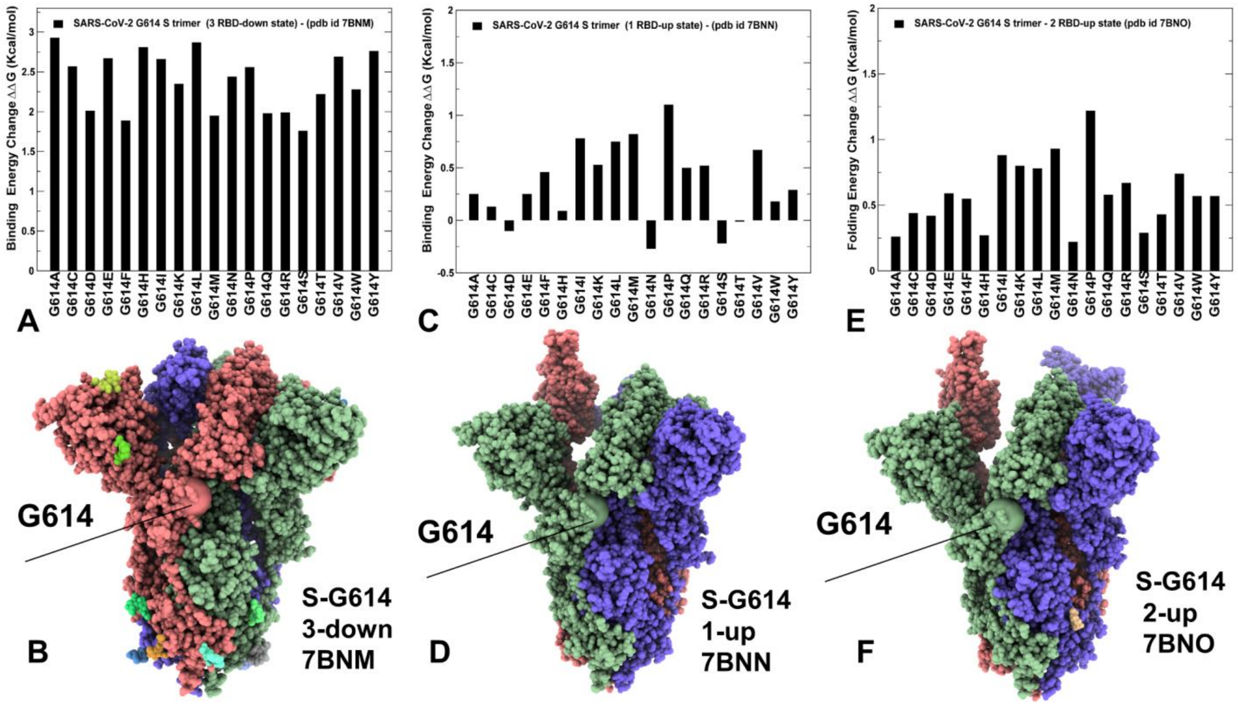


**Figure S4.** The mutational sensitivity analysis of the G614 residue in the SARS-CoV-2 S-G614 closed state, pdb id 7BNM (A,B), in the 1 RBD-up state, pdb id 7BNN (C,D) and 2 RBD-up open form, pdb id 7BNO (E,F). The structures are shown in full spheres and colored with protomers A,B,C are colored in green, red and blue. The position of G614 is shown in spheres.

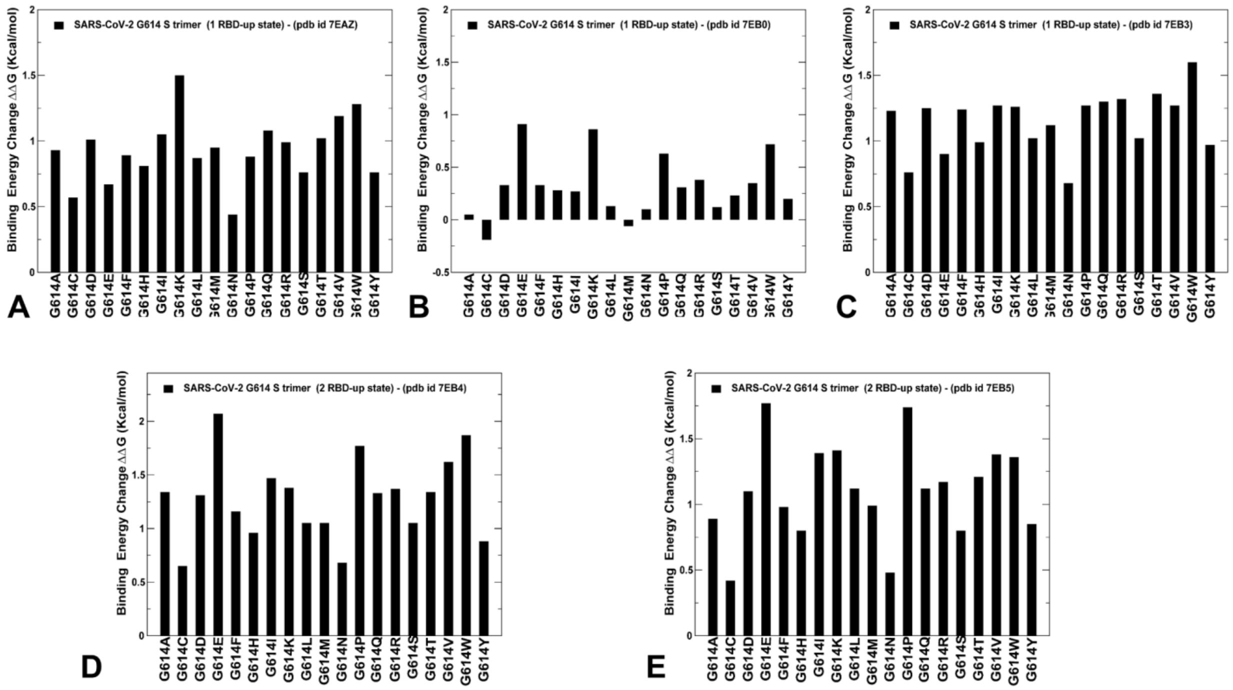


**Figure S5.** The mutational sensitivity analysis of the G614 residue in the SARS-CoV-2 S-G614 1 RBD-up open states (pdb id 7EAZ (A), pdb id 7EB0 (B), pdb OD 7EB3 (C)) and 2 RBD-up states (pdb id 7EB4 (D) and pdb id 7EB5 (E)).
